# Maintenance of parasite species diversity: Spatiotemporal niche partitioning and aggregation facilitate species coexistence

**DOI:** 10.1101/2024.10.21.619552

**Authors:** Ashwini Ramesh, Olivia McDermott-Sipe, Farrah Bashey-Visser

## Abstract

The maintenance of parasite diversity has historically taken a host-centric approach. Yet, many parasites are host generalists, and most parasites spend at least some time outside of their hosts. So, what mechanisms besides host-associated niche partitioning allow parasites to coexist? Using a year-long field survey and lab mesocosms, we examined whether environmental niche partitioning or aggregation could enable coexistence among soil-dwelling entomopathogenic nematodes. Field patterns along an elevational gradient reveal that species abundances differentiate with soil structure and moisture levels. Yet, most species strongly overlap within-sites throughout the year. Thus, niche partitioning alone is not sufficient to explain the coexistence of these species and other mechanisms are necessary to explain their coexistence. Aggregation at the within-site scale provides evidence for such a mechanism. Each species showed significant intraspecific clumping and largely random associations with other species. A mesocosm test of the consequences of intraspecific aggregation found that parasites at low or high densities limit their own population growth. Aggregation can promote negative feedback facilitating species coexistence. Our findings offer field-based evidence that spatiotemporal niche partitioning and aggregation both play a critical role in maintaining parasite species diversity, illustrating the importance of extending out view of parasites beyond their hosts.

## INTRODUCTION

Parasites are a critical, but often undervalued, component of all ecosystems (Hudson et al 2006). Loss of parasite species could destabilize food webs, thereby disrupt ecosystem function and even harm human health (Lafferty 2006, Torchin & Mitchell 2004). Thus, to best anticipate such harm, we need more fundamental insights into mechanisms favouring and undermining parasite species diversity. The maintenance of parasite diversity is predominantly viewed through the lens of the host (Hochberg & Holt 1990; Read & Taylor 2001). Parasites can specialize on different host species (Karvonen et al 2006) or use different strategies like competition vs. colonization or within-host niche partitioning to coexist on a single host (Bashey 2015; Ramesh & Hall 2024). While hosts are undeniably a crucial niche component, most parasites spend at least part of their life in a free-living stage outside the host, exposed to environmental conditions. These conditions can dramatically influence parasite survival (Carlson et al 2017) and their success in infecting new hosts (Handel et al 2014, Mideo et al 2009). This natural history of parasites raises a fundamental question: what ecological mechanisms maintain parasites species diversity outside of the host?

To answer this question, we can apply classic mechanisms maintaining species diversity from community ecology to the parasite realm. Spatiotemporal environmental niche partitioning is a well-studied mechanism of species coexistence (Fig. 1). Environmental variation can promote coexistence by allowing for niche differentiation (Brown et al 2013). When species differentiate their niches, they compete less for common resources, and thus are more likely to coexist (Albrecht & Gotelli 2001). In classic temporal niche partitioning, species adapted to different temporal conditions can lessen competition by being active at different times of the day or year (Fig. 1A; Hayward & Slotow 2009, Feau et al 2012). Similarly, in spatial niche partitioning, reduced spatial niche overlap can allow for multiple species to coexist across larger spatial scales, say, between sites or habitats (Fig. 1B; Chaudhary et al 2020, Wereszczuk & Zalewski 2015). Thus, spatiotemporal niche partitioning can allow species to coexist successfully over the course of a year across larger spatial scales.

**Figure 1:**
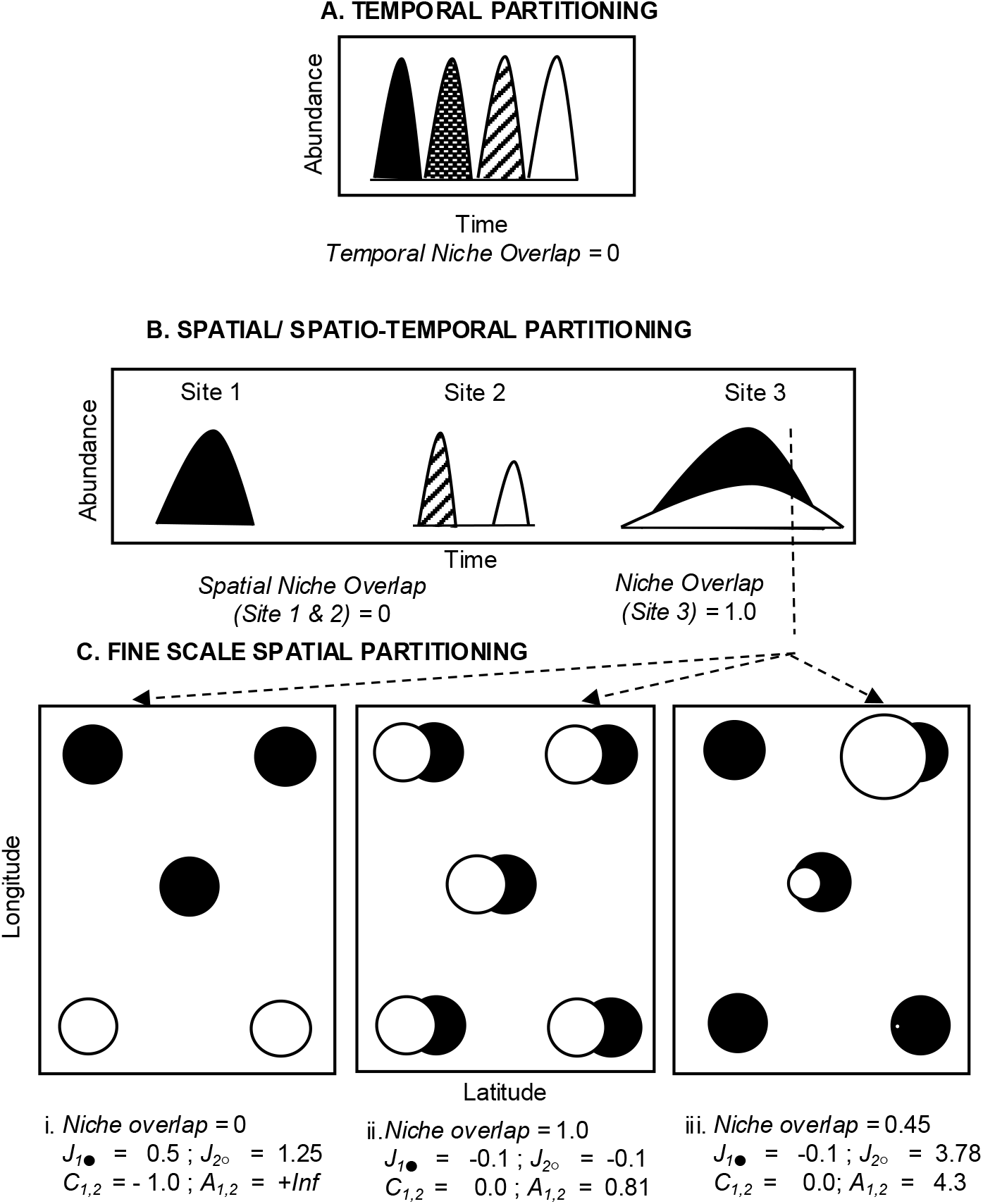
Niche partitioning across spatiotemporal scales: Species can coexist via classic niche partitioning at a variety of spatiotemporal scales. (A) *Temporal partitioning:* Species with similar niche requirements can lessen competition by being active at different times (e.g., between seasons) resulting in zero niche overlap allowing them to coexist. (B) *Spatial partitioning:* Species can reduce competition by occupying different sites (e.g., between Sites 1 and 2), or the same site at different times of the year under spatiotemporal partitioning (Site 2) facilitating coexistence (zero niche overlap). Species overlap can still allow for species coexistence, if niche partitioning occurs at a finer spatiotemporal scale, e.g., within a site at a given time (Site 3, time slice indicated by dashed line). (C) *Fine-scale spatial niche partitioning* facilitates coexistence if each species can occupy (i) different habitat patches within a site resulting in zero niche overlap. Typically, coexistence is not favourable when (ii) each species can occupy the same spatial patches within a site resulting in maximal niche overlap. However, if individuals of each species are overdispersed then aggregation can facilitate coexistence (iii). Aggregation is captured *via* variation in species distribution in abundance (circle size scaled to abundance). Intraspecific aggregation (*J*_*x*_) and interspecific aggregation (C_*x*,*y*_) determine species persistence (*A*_*x*,*y*_). Persistence requires that intraspecific aggregation is greater than interspecific aggregation. High intraspecific aggregation (*J*_*x*_ *> 0*) species can facilitate coexistence (*A*_*x*,*y*_ *> 1*) despite species occurring together in the same habitat patch (C_*x*,*y*_ >0). Here, we test these conceptual ideas of spatiotemporal mechanisms governing coexistence in four species of entomopathogenic nematodes at both (A and B) broader (between site and months) and (C) finer (within a site at a given time) scales.

However, what happens when species show strong spatiotemporal overlap at a given locale (e.g. as in Fig. 1B, Site 3)? Perhaps, niche partitioning can occur at a finer scale (Fig. 1C i). Species could specialize on specific microclimates or food resources (Luiselli 2006). Therefore, large-scale niche overlap maybe illusory, reflective of not being able to focus in on the key niche axes that differentiate species. Nevertheless, some species can show weak resource partitioning even at a local scale (Fig. 1C ii and iii, Takahashi et al 2005; Jaenike & James 1991). What then allows for species to coexist at finer spatial scales? Aggregation across an ephemeral or patchy niche, such as individuals of a host species, can allow for coexistence (Ives 1988, Sevenster 1996).

Aggregation refers to the degree to which individuals are clumped among habitat patches (Figure 1C). If most individuals of a given species occur in only a few patches, resulting in high variance in the number of individuals per patch, then the distribution is said to be intraspecifically aggregated. Analogously, interspecific aggregation is the degree to which two different species occur in the same patches, thereby increasing the positive covariance in the distributions of the two species. If intraspecific aggregation is greater than interspecific aggregation, individuals may compete more frequently with conspecifics than with heterospecifics, allowing both species to persist (Ives 1991). Importantly, intraspecific aggregation may reduce individual fitness such that each species imposes a brake on its own population growth. Several empirical studies have demonstrated the importance of aggregation in promoting coexistence in insect systems including *Drosophilia* (Sevenster & van Alphen 1996; Toda et al 1999), mosquitos (Fader & Juliano 2013), host-associated beetles (Ives 1991) and fleas (Krasnov et al 2006). In parasites, aggregated distributions are so prevalent that they are often regarded a universal law of parasite ecology (Poulin 2007a). However, this framework has seldom been used to test how aggregation supports parasite species coexistence (but see the preceding examples and Morand et al 1999) and has yet to be applied to environmental infective stages of parasites.

Aside from a few studies, research on the environmental distribution of parasite infective stages is generally lacking (Spiridonov et al 2007). Mechanistic studies of parasite distribution remain relatively rare, partly due to the significant logistical challenges they pose. Parasite distributions are typically described from a host-centric perspective using measures such as prevalence, mean intensity and even various indices of aggregation, sampled directly from the host. Yet, there is overwhelming evidence that environment distribution play a critical role in influencing infective stages even before host encounter, with significant implications for stabilizing host-parasite dynamics, hence community structure (Ettema 1998; Wilson et al. 2002). Despite these findings and ongoing calls for thorough investigation, substantial gaps in our knowledge of infective-stage distribution in the environment remain (Shaw & Dobson 1995; Morrill & Forbes 2016).

To address these challenges, here we examine mechanisms of species coexistence in a community of entomopathogenic nematodes (EPN). EPNs are soil-dwelling, insect parasites with a prolonged free-living juvenile stage (Strong *et al* 1996). EPNs can persist in soil for months, exposed to various environmental conditions before infecting a host. Although climatic factors like soil temperature and moisture significantly affect EPN dispersal, survival, and reproduction in lab settings (Bryant & Hallem 2018; Mason & Hominick 1995), linking these to field abundance remains difficult (Půža & Mráček 2005). Few studies have examined EPN coexistence in nature, and spatiotemporal niche partitioning has yet to be demonstrated (Půža & Mráček 2010). EPNs have broad host ranges and lack host specificity (Půža & Mráček 2010), making resource-based (i.e. host) niche partitioning unlikely. However, EPN distributions are typically aggregated, which has been hypothesized to contribute to their coexistence (Stuart & Gaugler 1994; Spiridonov et al 2007; Půža & Mráček 2010). Thus, the natural history of EPNs – survival in diverse environments, lack of host specificity, and aggregated distributions– make them ideal for investigating ecological mechanisms that maintain parasite species diversity beyond host niche axis.

The broad goal of our study is to test ecological mechanisms contributing to parasite species diversity at different spatiotemporal scales (Fig. 1). First, using a year-long field survey we establish the spatiotemporal distributions of four *Steinernema spp*. along a hillside gradient in a temperate forest. We examine the extent of species overlap in their distribution between seasons (Fig. 2), sites (Fig. 3) and how environmental gradients govern those species’ distributions. Second, we inspect the extent of species overlap in their distribution at the within-site scale (Fig. 4). Here, we test the role of aggregation in maintaining species persistence. Finally, using a lab mesocosm experiment we link how aggregation increases intraspecific competition causing each species to dampen their population growth (Fig. 5). Together, our study outlines how spatiotemporal niche partitioning and aggregation allow maintenance of nematode parasite species diversity.

**Figure 2:**
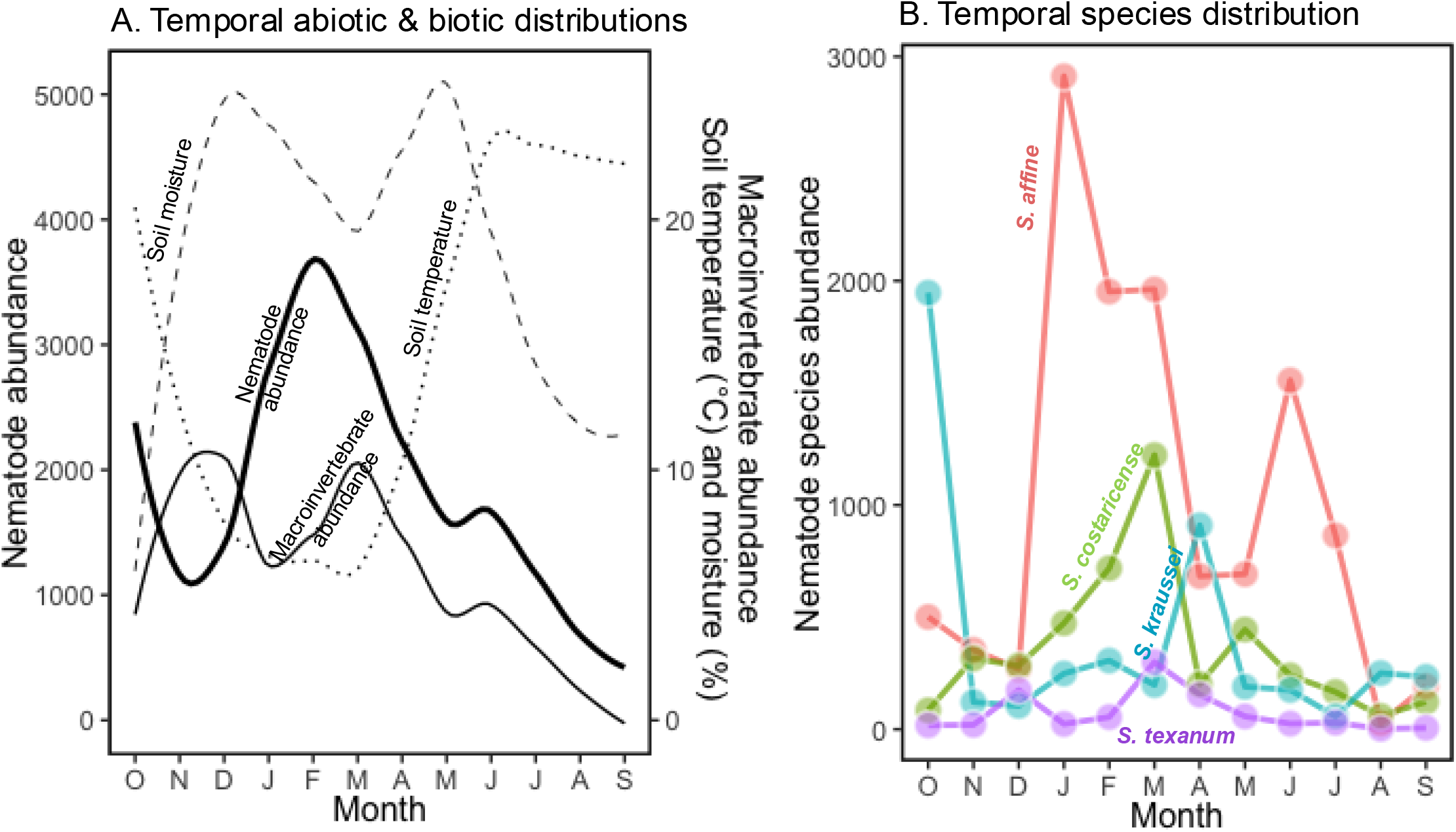
Temporal niche partitioning: (A) Overall, broad scale temporal distribution of nematode abundance (thick, solid line) summed over six sites, follows peaks in macroinvertebrate insect total abundance (thin, solid line) during cool temperature (^o^C, dotted line) and high soil moisture (% by volume, dashed line) conditions. (B) The nematode species abundance decomposed by four species of the genus *Steinernema*: *S. costaricense* (green), *S. affine* (red), *S. kraussei* (blue) and *S. texanum* (purple). Despite some variation in peak abundances, all species are present throughout the year, showing no significant niche partitioning by time.

**Figure 3:**
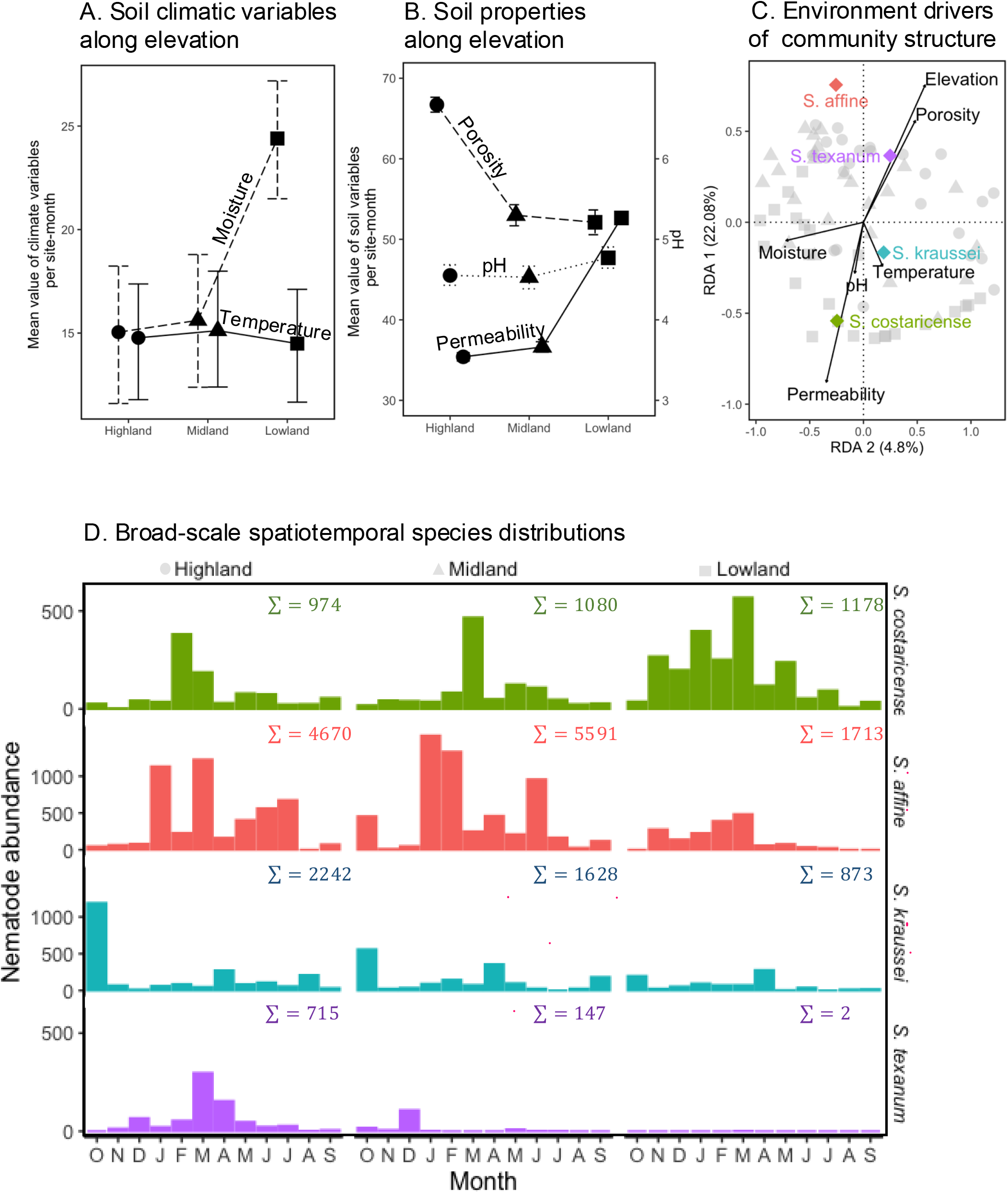
Broad scale spatiotemporal niche partitioning: Characterization of (A) soil moisture and temperature and (B) other soil properties along the elevational gradient (mean ±95% confidence intervals). (C) A redundancy analysis (RDA) examining the environmental effects (vectors) on nematode species abundances across each site-month combination (*n* = 72, labelled as highland ●, midlands ▲ and lowlands ◼) suggests that elevation explains the greatest variation in species composition. Each species centroid can be seen (coloured). (D) Monthly nematode species abundances along an elevational gradient; for each species, abundances are summed across two sites in each of the highland, midlands and lowlands. *S. costaricense* (green) abundances increase at lower elevations (lowlands), whereas *S. affine, S. kraussei* and *S. texanum* abundances decrease; with *S. texanum* largely absent in the lowland sites. Overall, while environmental gradients facilitate some niche partitioning, there was considerable spatial niche overlap at the broader scale.

**Figure 4:**
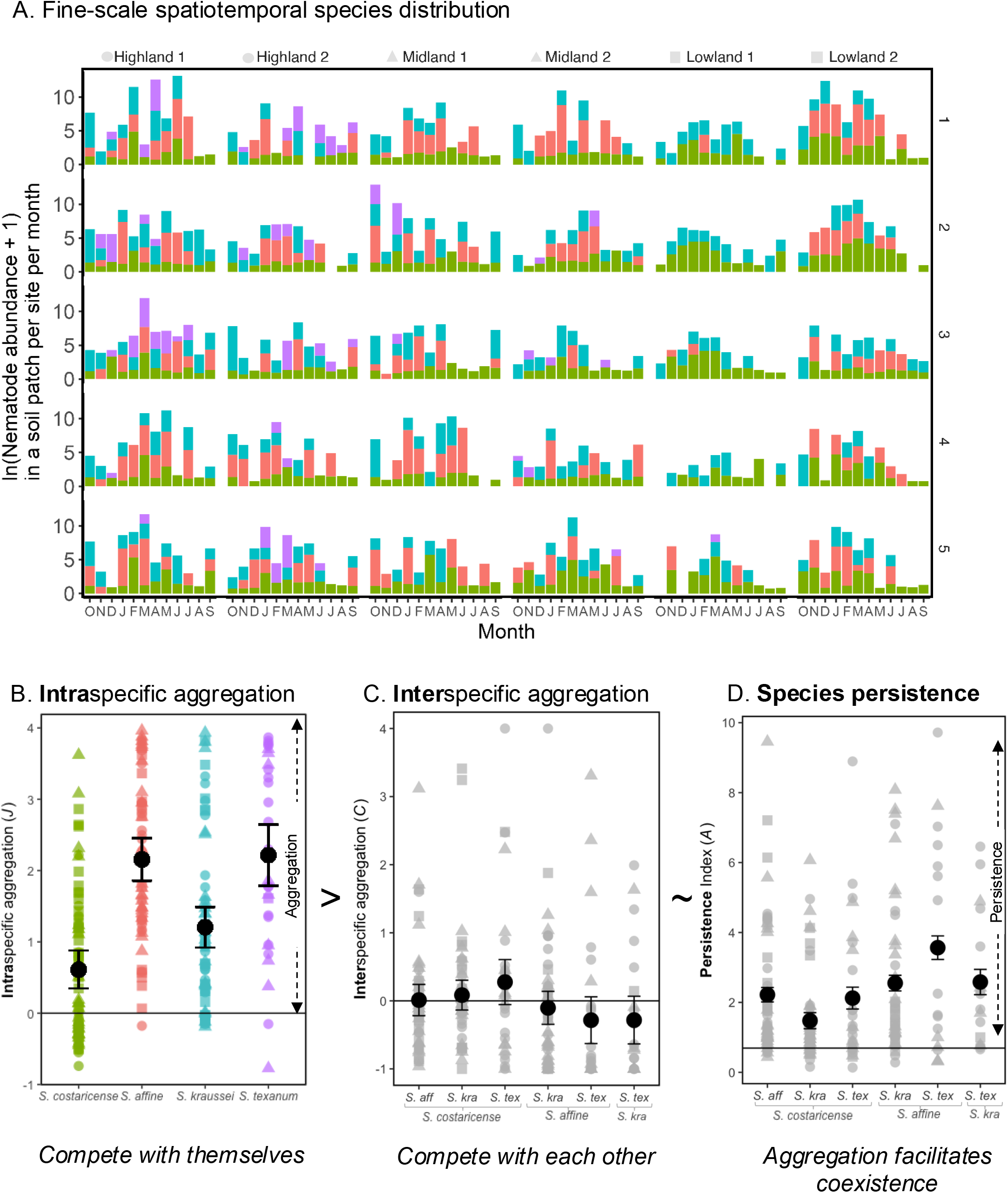
Fine scale spatial partitioning via aggregation: (A) Monthly measures of nematode abundances (left axis) from each soil sample taken from five distinct regions (right axis) within a site every month with *S. costaricense* (green), *S. affine* (red), *S. kraussei* (blue) and *S. texanum* (purple). Overall, species co-occur in 75% of the soil samples. (B) Intraspecific aggregation (*J*) indicates that all species are aggregated as the mean across all site-month combinations is significantly > 0, with *S. costaricense* significantly less aggregated (closer to random distribution) relative to other species. (C) Interspecific aggregation (*C*_*x*,*y*_), for each species pair *C*_*x*,*y*_ ∼ 0 indicates low or no association between species. (D) Together, the index of persistence (*A*_*x*,*y*_) which measures the relative effect of intra-*vs* inter-specific aggregation indicates that high intraspecific aggregation facilities species persistence (*A*_*x*,*y*_ *>* 1; here *A*_*x*,*y*_ is log transformed resulting in *ln*(*A*_*x*,*y*_ *+* 1) > 0.69, —). Points indicate indices calculated for each site-month combination, with highland ●, midlands ▲ and lowlands ◼. Large black dots are mean and error bars are 95 % confidence intervals.

**Figure 5:**
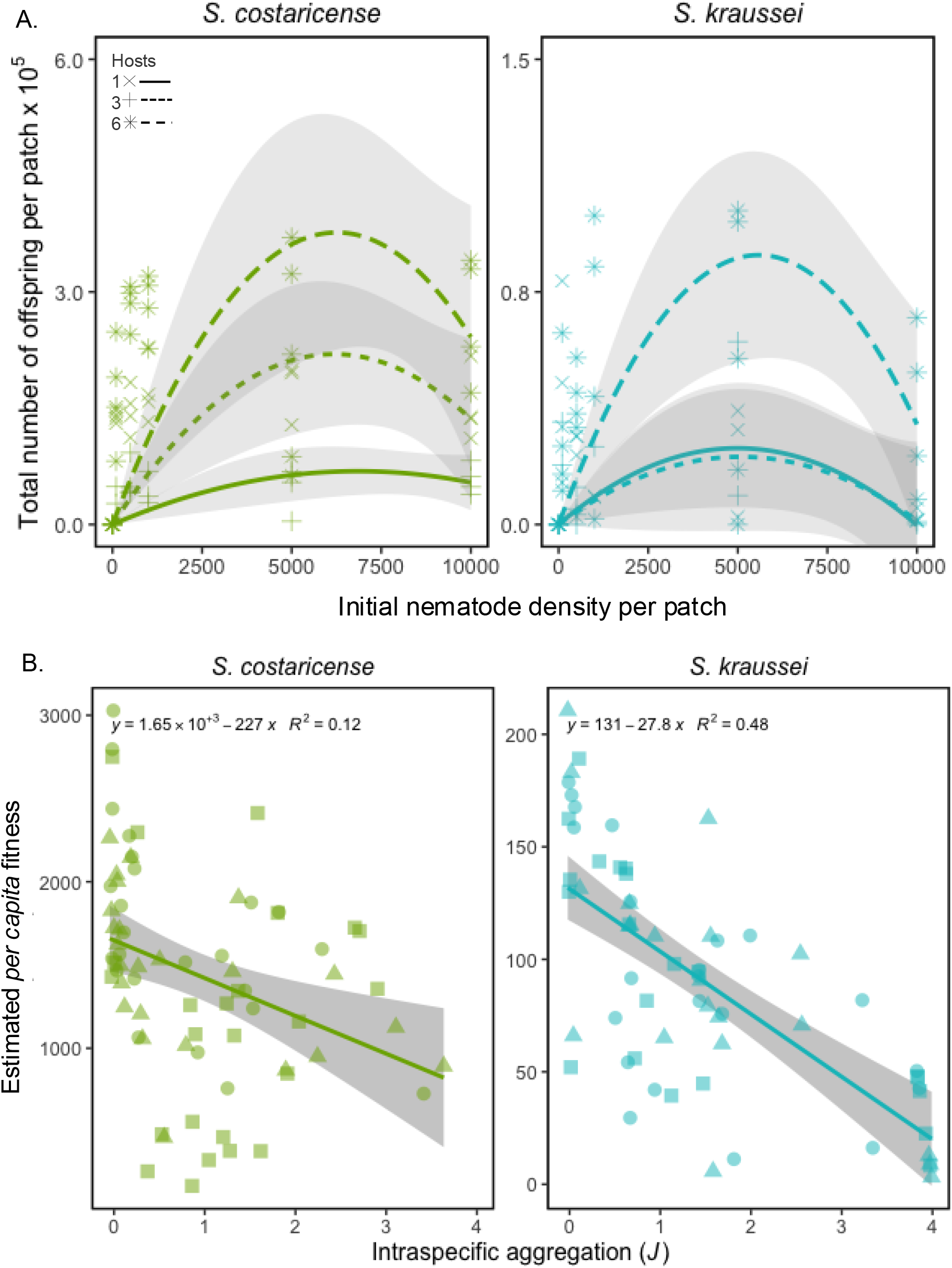
Intraspecific aggregation translates to intraspecific competition: (A) A lab mesocosm experiment measuring the effect of nematode density on total number of offspring produced per mesocosm across various host densities (one [solid]; three [small dashed]; six [long dashed] hosts) revealed a hump-shaped curve with total number of offspring highest at intermediate infecting densities for *S. costaricense* (green, left panels) and *S. kraussei* (blue, right panels). Aggregation results when individuals are found in relatively low or high densities, which lowers the total number of offspring produced by a patch relative to a more uniform or random distribution. (B) Estimated *per capita* nematode fitness calculated for each site-month from the field decreases with increasing intraspecific aggregation (*J*). Therefore, aggregation can dampen intraspecific growth allowing nematode species to coexist at finer spatial scales.

## METHOD

### Study System

Nematodes in the genus *Steinernema* are entomopathogenic with a prolonged free-living juvenile stage. The soil-dwelling juveniles are non-feeding and developmentally dormant but can actively seek insect hosts (Bal & Grewal 2015). EPNs are considered host generalists as they can infect and have been isolated from a wide variety of soft-bodied insects (Kaya & Gaugler 1993, Peters 1996). On entering an insect host, nematodes resume their development, quickly kill the host, forage, mate, and reproduce within the carcass (Selvan *et al* 1993). Offspring emerge from the host in 3-4 weeks and continue the cycle. EPNs can persist in soil for several months where they are exposed to a variety of environmental conditions like varying temperature, moisture, pH, before they successfully infect a host. The limited host availability can restrict the proliferation and spread of EPNs (Půža & Mráček 2010), thereby contributing to their spatial patchiness and niche partitioning in the environment (Stuart & Gaugler 1994).

### Study site and sampling

We examined nematode communities in Moores Creek, Indiana a temperate forest reserve located in the Indiana University Research and Teaching Preserve, Bloomington, IN, USA (39°05′ N, 86°28′ W) built on the lands of myaamiaki, Lënape, Bodwéwadmik, and saawanwa people. From October 2019 to September 2020, we collected monthly soil samples across six forest sites (4 x 4 m^2^). The sites were along a hillside gradient, with two sites each (60 meters apart) at the high (elevation = 218 m; Highlands), middle (216-212 m; Midlands), and lowest end (205-208 m; Lowlands) of the gradient, with elevations extracted using the GeoNames geographical database (Rowlingson 2024). Each site was divided into five regions, and every month a unique soil sample (∼300 cm^3^) was taken from the topsoil of each region using a 4 cm diameter by 8 cm deep soil corer. To isolate the potential host organisms, soil samples were gently sifted by hand. All macroinvertebrates (over 3 mm in body length) were preserved in 70% ethanol and identified to the genus level. To isolate nematodes, soil samples suspended in one litre of water were manually filtered through a 2000 *μM* and 20 *μM* mesh sieves (Hogenogler & co). The filtered residue was further processed using a sucrose centrifugation method (Freckman & Viginina 1997). DNA was then extracted from each sample using the Qiagen DNeasy PowerSoil Pro Kit. From the resulting mixed DNA sample, we quantified the abundance of each the four species of the genus *Steinernema* present at the study site: *S. affine, S. costaricense, S. kraussei* and *S. texanum*, using species-specific primers in a standard qPCR (S1). Thus, we characterized the temporal and spatial dynamics of the nematode community using 360 unique samples (12 months x 6 sites x 5 soil samples).

To examine the role of environmental variables driving nematode community dynamics, we measured two broad classes: variables fluctuating with seasonal changes, such as temperature and moisture, and those spatially fixed, including soil pH, porosity, and permeability. Soil temperature and moisture (volumetric water content, VWC) were measured using handheld sensors (HydroSense II, Campbell Scientific) in each of the soil cores (or patches) in the field per site per month. Soil pH was measured in each of the soil samples in the lab after first suspending 5g of dried soil sample in 0.01M CaCl_2_ (Soil S.P.A.C, 2000). Soil porosity is the ratio of the volume of openings (pores) to the total volume of material, while permeability is the speed at which water moves through interconnected pores within soil. We collected soil samples separately after the twelve-month survey, given that soil structure was disrupted during the sieving process and soil properties typically do not change across season. To measure soil permeability and porosity, we sampled roughly sixteen soil samples that spanned each of the sites for total of 92 samples. Soil porosity was estimated by measuring the bulk and particle densities (Blake & Hartge 1986), while soil permeability was determined by the time taken for 100 mL of water to pass through the dry soil and its interconnected pores within (Klute & Dirksen 1986). The soil permeability and porosity were mapped onto the field dataset by using the average value of three values taken for each five regions sampled at each of the six sites. All statistics and visualizations were using R (version 4.0.5) (R Core Team, 2020).

## Evaluating broad scale temporal and spatial niche partitioning

To examine whether temporal and/or spatial niche partitioning facilitates species coexistence across our 300 m hillside gradient, we first examined the broad scale temporal pattern across the year and then examined the spatial pattern along the gradient.

### 1. Temporal niche partitioning

Can temporal partitioning facilitate species coexistence? First, we examine how seasonal factors like soil temperature and moisture affected nematode and potential host distributions. Seasonal factors were averaged across all sites and soil samples for each of the 12 sampled months, while nematode and macroinvertebrate abundances were summed across all sites and soil samples (*n* = 30, 6 sites x 5 soil samples) for sampled months. If species are adapted to different temporal conditions, reducing competition by being active at different times of the day or year, then temporal niche partitioning can promote species coexistence.

To quantify niche partitioning between species, we calculated Pianka’s index of niche overlap between each pair of species (Pianka 1973) using EcoSimR 1.00 (Gotelli et al 2015):

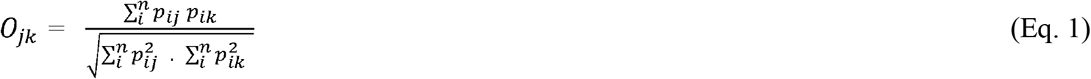

where *O*_*jk*_ is Pianka’s index of niche overlap between species *j* and *k, p*_*ij*_ is the proportion of the *i*th resource used by species *j, p*_*ik*_ is the proportion of the *i*th resource used by species *k*, and *n* is the total number of resources. Here, the resource is the habitat (sampling sites or time), the use of which is assumed to be expressed by species abundances (Tsafack *et al* 2021). The *O*_*jk*_ index is symmetric where values close to 0 reflect usage of exclusive resource categories, whereas values close to 1 reflect similar resource utilization spectra. The statistical significance of the niche overlap patterns across the four species in our assemblage was determined by comparing the observed mean of all possible pairwise species overlap indices to a distribution based on randomized resampling of the species usage data. Specifically, we used the RA4 algorithm where only the non-zero entries for each of the four species were randomly reshuffled across resource habitats (here the 12 months). This procedure retains both the niche breadth of the species and the pattern of zero states (Winemiller & Pianka 1990, Albrecht & Gotelli 2000). We tested whether there was significant niche partitioning by examining the lower 95% confidence limit based on a one-tailed test from 1000 simulated usage matrices. To calculate temporal niche overlap, each row of the data matrix represented species abundances summed across all sites (*r = 4*) and each column represented a month of the year (*n* = 12 months).

### 2. Spatial niche partitioning

To test if spatial niche partitioning facilitated species coexistence, the changes in environmental variables at each site along the hillside were examined. A mixed-model ANOVA tested whether soil temperature, moisture, and pH varied across sites based on the average within-site soil samples (*n* = 72: 6 sites x 12 months) collected during the survey with month used as a random effect. Soil porosity and permeability variation by site were assessed with an ANOVA based on the 92 soil samples collected after the monthly survey. Nematode abundances were then examined to evaluate whether species differentiate their spatial niches. Patterns of nematode abundance were visualized by pooling data gathered each month across two neighbouring sites along the elevational gradient. The spatial difference in species abundances was tested using a mixed effects model with a negative binomial distribution, with elevational categories as fixed effects and month as a random effect (Bates et al 2015). To test the extent of niche overlap, each row of the data matrix represented species abundances averaged across the year (*r* = 4 species) and each column represented the six sites (*n* = 6 sites).

### 3. Spatiotemporal niche partitioning

The variation of nematode abundances across site-month combinations was further examined using a multivariate analysis. A principal component analysis (PCA) on Hellinger-transformed abundances visualized spatial patterns of community composition for every site per month (*n* = 72: 6 sites x 12 months) (Legendre & Legendre 1983). Redundancy analysis (RDA) was used to identify environmental drivers of community dynamics. Variation partitioning via RDA showed that abiotic environmental variables (elevation, permeability, porosity, pH, temperature, and moisture) explained 18.6% of the variation in nematode species composition, while macroinvertebrate abundance explained less than 1%. Given this, we present the results of an RDA examining the effect of abiotic environmental variation on nematode abundance. The combination of space and time’s effect on niche overlap of the nematode species was examined. The niche overlap index was calculated using a data matrix where each row represented summed species abundances for each site in a given month (*r* = 4) and each column represented the (*n* = 72 = 6 sites x 12 months).

### Fine scale spatiotemporal niche partitioning

If species overlap at larger spatiotemporal scales, coexistence may be possible if niches are partitioned at finer scales. Thus, the abundance of each nematode species in each of the 360 soil samples was examined to calculate patterns of co-occurrence. Additionally, niche overlap of the four species across these 360 soil samples was tested. Finally, niche overlap between pairs of species within each site for every site-month combination was calculated.

#### A test of aggregation model of coexistence

To test if aggregation contributed to patterns of coexistence, we tested the relative effect of intra- and inter-specific aggregation in maintaining species persistence. Aggregation was measured at the within-site scale across the five soil samples collected every month. Here, let *n*_*i*_ denote the number of individuals of species x in a patch *i* i.e., unique soil sample, *m*_*x*_ the mean number of individuals per patch, and *N* the total number of individuals of species *x* in a site at that time. If individuals were randomly and independently distributed among patches, they would follow a Poisson distribution such that the expected value of 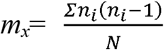, (Ives 1988). If species is aggregated the right-hand side will be greater than *m*_*x*_. Therefore, the degree of intra-specific aggregation is expressed by the index *J* is the proportional increase in the observed density of conspecific competitors relative to a random distribution (i.e., a Poisson distribution) expressed as:

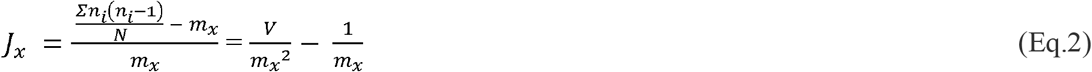

If *J*_*x*_ = 0, then variance (*V*) is equal to the mean and species are randomly distributed, *J*_*x*_ > 0 implies variance much greater than the mean i.e., over-dispersion or aggregation, and *J*_*x*_ < 0 is under-dispersion. High intraspecific aggregation is stabilizing.

Interspecific aggregation between two species is measured using an analogous index: *C*_*xy*_

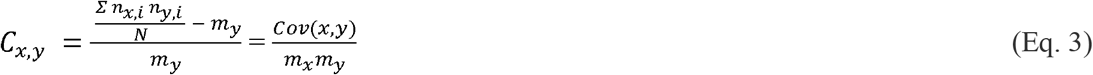

where *cov(x*,*y)* the covariance between species *x* and *y*. If *C* = 0, then there is random association between the species, or low interspecific competition. If *C* > 0, there is positive association implying high potential for interspecific competition; *C* < 0 there is negative association (or no overlap) indicates resource segregation.

*J*_*x*_ and *C*_*x*,*y*_ can be combined to obtain a condition for persistence. Under reasonable assumptions (see Sevenster 1996), species persistence (*A*) is defined as:

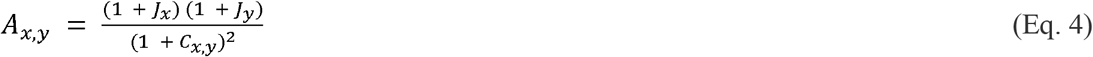

*A*_*xy*_ >1 is a necessary condition for the persistence of a focal species *x* in the presence of species *y* and vice-versa. This occurs when the joint intraspecific aggregation of each species is greater than their interspecific aggregation, indicating stabilization. When *A*_*xy*_ ⊔ 1 then then each species is unlikely to be able increase from low density and persist in the presence of its competitor.

Here, we analyse species distributions, niche overlap, and their impact on persistence for each site-month combination. Figure 1C provide an illustration for three representative cases, where species abundance (proportional to the size of circles) for a species pair (black and white) is measured across five soil samples (patches) taken within a site at a given month. In the first case (Fig. 1C.i), if both species, species 1 (●) and species 2 (○), strictly partition their niches (i.e., Niche overlap = 0), then competition (*C* _●,○_ = −1) facilitate species persistence (*A* _●,○_ > 1). This represents the classic case of resource partitioning. In the remaining two cases, (Fig. 1C.ii-iii), species ● is spread nearly evenly across all patches as captured by its intraspecific aggregation, *J*_●_ *∼ 0*. Now, if species ○ is also evenly distributed, or has low intraspecific aggregation (*J*_○_ ∼ 0, Fig. 1C.ii), then the randomly association between the two species (*C* _●,○_ ∼ 0) allows neither to persist (*A*_●,○_ < 1). However, if species ○ is clumped or highly aggregated (*J*_○_ > 0, Fig. 1C.iii) then individuals compete more frequently with themselves than with their competitors allowing both species to persist (*A*_●,○_ > 1). Thus, aggregation rescues species from competitive exclusion facilitating coexistence.

To identify if the aggregations indices were significantly different from zero for *J*_*x*_ and *C*_*x*,*y*_, and whether they differed among species, we ran a generalized linear model for each index with species as a fixed effect and site and time as random effects. We conducted a similar analysis for *A*_*xy*_. To address skewness, we log-transformed *A*_*xy*_ and applied the glm.nb function from the MASS package (Venables & Ripley 2002) to implement a generalized linear model with a negative binomial distribution. This allowed us to test whether *A*_*xy*_ > 1 or (*A*_*xy*_ + 1 > *ln*(2) = 0.69. We then used the 95% confidence intervals to determine if the estimates fell within the range of their respective thresholds

### Mesocosm Experiment

#### Does increasing intraspecific aggregation imply intraspecific competition?

Coexistence via aggregation assumes that intra-specific aggregation will increase intraspecific competition relative to interspecific competition, such that each species dampens its own population growth. To investigate if aggregation contributes to stabilization through intraspecific competition: (1) we analyzed the relationship between nematode density and per-patch productivity in lab mesocosms to estimate per capita fitness from field soil samples; and then (2) show that how this density-fitness relationships results in a negative correlated between fitness and intraspecific aggregation (*J*) across site-month combinations.

First, to establish nematode density-fitness relationship we conducted a lab mesocosm experiments. Soil-filled garden pots (4”d x 4”h) were prepared using ∼350 cm^3^ of 50:50 mixture of local soil and Metro Mix (Sun Grow Horticulture), twice autoclaved. Each pot was seeded with one of seven initial nematode densities (0, 100, 500, 1000, 5000, 10000) and one, three, or six potential hosts, larvae of wax moth, *Galleria mellonella* (Vanderhorst Wholesalers). Here, we focussed on two species: *S. costaricense* and *S. kraussei*. Caterpillar mortality was assayed within seven □ days post-infection; dead caterpillars were moved individually to modified White traps to allow for the emergence and collection of juvenile nematodes (Bashey & Lively, 2009). Both mesocosm and white traps were kept at 20°C and soil moisture in the mesocosm was 20% VWC. Caterpillars were assayed for emergence and infective juvenile nematodes were collected until six □ weeks post-infection. The number of nematode offspring was estimated by volumetric subsampling. We used a generalized linear model with a quadratic relationship to determine how the initial density of nematodes in the mesocosm affected the total number of offspring produced per mesocosm.

Based on the size of our soil samples and macroinvertebrate abundance, nematodes found in each of the field soil patches are likely to compete at most for single host. Consequently, we used the quadratic relationship between total number of offspring and initial density for a single host mesocosm to estimate the *per capita* fitness 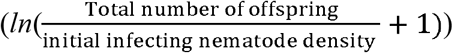 for each field soil sample. █Earlier, we presented nematode abundance estimates based on direct qPCR data for all species. For fitness quantification, we limited our analysis to species that could be readily cultured in the lab, adjusting for losses during sieving to achieve more precise abundance estimates. We sieved known quantities, performed qPCR, and backtransformed the data to calculate per capita fitness per soil sample, leading to the mean fitness estimate for each site (S3.2). Using these fitness estimates and the more precise density estimates, we investigated how site-month nematode fitness related to intraspecific aggregation at each site. To assess the impact of aggregation on parasite fitness, we regressed the mean per capita fitness against intraspecific aggregation (*Jx*).

## RESULTS

### Evaluating broad scale temporal and spatial niche partitioning

#### 1. Temporal niche partitioning

Across the study site, the highest peak in nematode abundance followed the abundance of potential invertebrate hosts (Fig. 2A). The most prevalent invertebrates were beetle and fly larvae (TableS2). Nematodes were most abundant in the late winter, i.e. a period of low temperature and high moisture. Smaller peaks in nematode abundance were observed in hotter, drier months (Fig 2A). This general pattern was mainly driven by the most abundant species, *S. affine*, which peaked in January and again in June (Fig 2B). While the next most abundant species, *S. kraussei*, peaked in October and in April. Both *S. costaricense* and *S. texanum* peaked in March.

Despite these distinct abundance peaks, all species were present throughout the year, overlapping in their temporal niche. The species showed 54.9% niche overlap based on their monthly abundances, meaning they were closer to showing similar patterns in their usage of time than to having distinct temporal profiles. This degree of overlap was not significantly less than the null expectation based on independent temporal habitat usage across species (null = 47.5%, *p* =0.88). Thus, species distribution across months provide insufficient support for temporal niche partitioning as a mechanism to explain species coexistence.

#### 2. Spatial niche partitioning

If not in time, do species separate out in space instead? Our study sites were along a mild elevation gradient. Moving down the gradient, significant changes were observed in several environment variables (Table S5A). Most distinct were the soil samples from the lowest elevation sites, which had significantly higher moisture content, permeability, and pH, but lower porosity (Figure 3A and B). Additionally, while soil temperature was mostly affected by season, we found the mid-elevation sites had a slight, but significant increase in temperature, most likely due to their south-facing aspect.

The patterns of nematode abundance can best be visualized by pooling nematode abundance data gathered each month across two neighbouring sites along the elevational gradient (Fig. 3D?). Overall, species show distinct distributions. The total abundance (Σ in panel inset) of *S. costaricense* increased down the hillside gradient with abundances significantly higher in the lowlands (*F*_*2*,*67*_ = 8.0, *p* < 0.001). In contrast, *S. affine* and *S. kraussei* abundances were significantly lower in the lowlands (*S. affine*: *F*_*2*,*67*_ = 2.58, *p* < 0.001, *S. kraussei F*_*2*,*67*_ = 3.08, *p* < 0.001,). Notably, *S. texanum* was mostly found in the highlands and virtually absent in the lowlands (*p* < 0.001, *F*_*2*,*67*_ = 904; Table S5C).

Despite these different patterns, species showed an even higher degree of niche overlap (72.4%) when examined across the six sites than seen temporally. This degree of overlap was not significantly less than the null expectation based on independent site usage across species (*null* = 70.0, *p* = 0.7094). Therefore, across sites, spatial niche partitioning was insufficient to explain species coexistence.

#### 3. Spatiotemporal niche partitioning

We further examined how nematode abundances varied across site-month combinations by using a multivariate analysis. Environmental variables explained 24.2% of the variation in nematode species abundances across the 72 site-month combinations. In the RDA triplot (Fig. 3C), a species abundance is maximal near the species centroid (⍰) and decreases away from it. Thus, site points (highland ●, midlands ▲ and lowlands ◼) near the species centroid reflect the spatial niche separation. The longer the black arrows corresponding to an environment factor are, the more strongly it is associated with variation in nematode species abundance matrix. Of the canonical axes, which represent variation in explanatory variables in fewer dimensions, only RDA 1 was statistically significant (*p* = 0.001), explaining 22% of the variation, while the RDA 2 explained 4.8%. The full model revealed that elevation and permeability load strongly on the first axis (*p* < 0.01), while moisture loads more strongly on the second axis. Porosity, pH, and temperature also load on the first axis but to a lesser extent (S2.2 Table S4).

The relative positioning of environmental variables and species centroids in the RDA plot indicates the effects of the environmental gradients on the species relative abundance (Fig. 3C). *S. texanum*’s centroid lies close to the vectors for elevation and porosity, indicating their positive effects on its abundance. *S. texanum* is restricted to the upper sites and nearly absent in the lowlands (Fig. 3D). Porosity, which was significantly higher in the highlands (Fig. 3B), refers to the space between soil particles. These spaces in the drier upper soils may provide key habitat for *S. texanum*. In contrast, *S. costaricense*’s centroid is found in the opposite (lower left) quadrant of the RDA plot (Fig. 3C), indicating its negative association with porosity and elevation, and the positive effect of permeability and pH on its abundances. Higher permeability means that water passes through soils quickly, and the placement of *S. affine*’s centroid in the opposite (top left) quadrant, suggests that *S. affine* may do better with soils that retain their moisture longer and are more acidic than those where *S. costaricense* showed higher abundance. Finally for *S. kraussei*, the major environmental predictor is temperature, reflecting its abundance peaks during the relatively warm month of October.

Thus, the multivariate analysis suggests that the species do differ in their usage of the environment. In fact, the niche overlap across the 72 site-month combinations was decreased to 22.5%, which is less than, but not significantly different from, the null expectation (null = 24.1, *p* = 0.18). Thus, nematodes show more distinct usage patterns across sites and months, but this separation was not sufficient to be regarded as significant spatiotemporal niche partitioning.

### Fine scale spatiotemporal niche partitioning

If species overlap at broad spatial and temporal scales, do they instead separate out at at finer spatial scales? To test such fine scale spatiotemporal niche partitioning, we first examine species distributions in each soil sample (Fig 4A). Out of the 360 soil samples, 351 contained nematodes. Of these, nematodes were found to co-occur as species pairs in 42% of the samples, in triads in 31% of the samples, and even in tetrads (3%). In fact, nematode species were only found in isolation in less than one quarter of the samples. However, when examining the relative abundances of each species, species show niche partitioning at this finer spatial scale. Calculating the niche overlap across the 360 soil samples, shows that this is decreased to 5.9%, which is lower than the null expectation of 7.7% and approaching significance (one-tailed *p* = 0.05).

Nevertheless, much of the differences in soil sample useage as seen in Figure 4A appears to be reflected in the broader spatiotemporal patterns already seen in Figure 3B&C. Thus, we calculated niche overlap at the within-site level for each of the 72 site-month combinations by using the relative abundance of each species in the five soil samples. Only 2 of 72 site-month combinations showed significant niche partitioning. For example, in June at the Midland-2 site, even with *S. costaricense* (green) present in every soil sample, the other three species occupied different soil samples, demonstrating niche partitioning. The remaining site-month combinations showed no significant partitioning with the mean overlap 40%. (S2.1 Fig S2). Therefore, at the within-site scale species do not show spatial niche partitioning, but mostly co-occur within a patch suggesting that some other mechanism must maintain species coexistence across patches.

### A test of aggregation model of coexistence: Do co-occurrence patterns favour coexistence?

The index of persistence examines the degree to which self-aggregation and niche partitioning can permit each species to persist in the community. Here, we analyse species distributions, niche overlap, and their impact on persistence for each site-month combination (Fig. 4B-D; Table S6).

The intraspecific aggregation index *(J)* compares the distribution of abundances across patches to a random distribution, such that *J* > 0 indicates aggregation. In general, all species are aggregated, showing a mean aggregation significantly greater than 0 (Fig. 4B, Table S6). This aggregation may result in self-dampening for these species. Among the species, *S. costaricense* is significantly less aggregated compared to the other species (*F*_*3*,*218*_ = 24.56, *p* < 0.01). In fact, 31/72 site-month combinations show uniform distribution across patches (i.e. *J* < 0) for *S. costaricense*, while only 15/72 for the remaining species.

The index of interspecific aggregation *C*_*x*,*y*_ was centered around zero for all species, indicating that there was random association between the species (Fig. 4C, Table S6). Thus, while there is the potential for interspecific competition within a site at a given time, it is predicted to be relatively weak compared to interspecific competition. Importantly, average *C*_*x*,*y*_ is close to zero and not negative, indicating that resource partitioning or competitive exclusion is not occurring at this scale

Taken together, the index of persistence (*A*_*x*,*y*_) which measures the joint effect of aggregation of each species on its competitor and vice-versa predicts persistence for most species pairs (*A*_*x*,*y*_ > 1 or *ln*(*A*_*x*,*y*_ + 1) > 0.69; black line; Fig. 4D,Table S6). Therefore, nematode distributions of *Steinernema* spp. were each sufficiently aggregated (*J > 0*) to permit the continued persistence of nearly all species pairs within each site presently occupied.

### Mesocosm Experiment

#### Increasing intraspecific aggregation increases intraspecific competition

Species persistence (*A*_*x*,*y*_) is driven by high intraspecific aggregation (*J*) rather than negative interspecific aggregation (*C*_*x*,*y*_; Fig. 4B-D). Does this high within species aggregation translate to high intraspecific competition? Lab mesocosms with varying initial densities were used to test the fitness consequences of intraspecific aggregation for two of the four species: *S. costaricense* and *S. kraussei*. A family of quadratic curves governed the relationship between number of infecting juveniles in the soil and total number of offspring per patch, across all combinations of host densities in both species (Fig. 5A; Table S7). Specifically, too few or too many nematodes occupying a mesocosm reduced number of offspring, but intermediate densities increased offspring numbers.

Depending on nematode distribution at the within-site scale, the non-linear consequences of nematode density on offspring production per patch can vary the degree of negative feedback at the population scale. Given the low abundance of potential insect hosts observed in the field survey, we assume that nematodes in a given patch of soil will compete for access to a single host. To capture the fitness effect of aggregation observed at the within-site scale in the field, we calculated the *per capita* nematode fitness per site month-combination based on the observed nematode abundances in each soil sample. Then, we examined how within site-level fitness related to intraspecific aggregation (*J*) observed for each site-month. As expected under the quadratic relationship between initial nematode density and total number of offspring, increasing aggregation reduced *per capita* fitness per patch for both species (*S. costaricense*: *R*^2^ = 0.12, *p* < 0.05; *S. kraussei*: *R*^2^ = 0.48, *p* < 0.05; Fig. 5B). This suggests that the key assumption of the aggregation framework, that that greater intraspecific aggregation leads to increased intraspecific competition, is true for *Steinernema spp*.

## DISCUSSION

What ecological mechanisms maintain parasite species diversity? Here, we investigated this question by testing three competing classic niche-based mechanisms of species coexistence. During infective stages, species can coexist if they successfully differentiate their niches via (i) temporal niche partitioning (ii) spatial niche partitioning or (iii) aggregated distributions that result in intraspecific dampening (Fig. 1). First, using a year-long field survey, we examined whether spatiotemporal environmental niche partitioning among soil-dwelling entomopathogenic nematodes could explain their coexistence. At the larger spatiotemporal scale *i*.*e*. between sites sampled monthly, environmental gradients favour species in some sites but nearly exclude them in others (Fig. 2-3). Yet, most species strongly overlapped across seasons and sites, limiting the role of spatial or temporal niche partitioning in maintaining coexistence. Species distributions at a finer spatiotemporal scale, *i*.*e*. within sites for each month in the year, revealed that species overlap still (Fig. 4). However, most species were over-dispersed or aggregated with themselves but randomly associated with their competitors, leading to greater intra-relative to inter-specific aggregation. A lab mesocosm experiment then demonstrated that when nematodes are aggregated in a patch, nematode fitness was reduced, due to the non-linear relationship between the number of nematodes in a patch and the total number of offspring produced (Fig. 5a). Intraspecific aggregation increases intraspecific competition and lowers intraspecific facilitation, creating a negative fitness feedback (Fig. 5 b) which can promote species coexistence. Our study is among the first to link field and lab measurements of environmental distributions of parasite infective stages to determine the mechanisms underlying species coexistence. Overall, by examining the mechanisms of parasite coexistence across scales, we demonstrate how environmental niche partitioning and coexistence via aggregation both facilitate the maintenance of parasite species diversity.

### Broad temporal and spatial scales: Coexistence via environment niche partitioning

Environmental niche partitioning favoured species in some sites but exclude them in others. Yet, most species strongly overlapped across seasons and sites, limiting the role of spatial or temporal niche partitioning in maintaining coexistence. All species were present monthly throughout the year (Fig. 2B) and showed strong temporal niche overlap. Similarly, species abundances across six sites along an elevation gradient also showed strong niche overlap, with three of the four species occurring in all sites (Fig. 4A). A hint of niche partitioning was found when examining species relative abundances across 72 site-month combinations. While not significant using a randomization test of Pianka’s niche overlap index, species were found to respond differently across the gradient, suggesting that differential responses to environmental variation could facilitate species coexistence (Fig. 3C). Most noticeably, *S. costaricense* increased in abundance in the in the lowlands, while the other species decreased, with *S. texanum* being virtually absent (Fig. 3D). While highland and midland sites were characterized by similar levels of soil moisture and permeability, the lowland sites exhibited a 1.5-fold increase in these parameters (Fig 3A and B). The intensified soil moisture in lowlands possibly limits the survival of the other species, but favours *S. costaricense*. In contrast, the highland sites stand out with increased soil porosity (Fig 3B), these spaces may provide key habitat for *S. texanum* in the highlands. Soil properties can differentially effect infectivity and survival of EPNs (Erktan et al 2020, Koppenhoeferi et al 2007, Matuska-Lyzwa et al. 2024), but these effects have yet to be characterized for the species we found in this forest community.

Environmental tolerances shape parasite niche breadth outside the host facilitating a parasite-centric approach to understanding parasite diversity. Historically, those tolerances have been studied in ectoparasites due to the applied interest in controlling the diseases they harbor (Laporta et al. 2014). For instance, extensive geo-referenced data reveals environmental niche overlap among tick species, their hosts, and pathogens (Estrada-Pena & de la Fuente 2016). Similarly, many non-feeding macroparasites actively move to infect hosts, relying on narrow time and temperature windows for effective transmission (Diaz-Munoz et al. 2023). These selection pressures also extend to other deadly diseases; for example, avian influenza strains demonstrate a trade-off between persistence at low temperatures and resilience to high temperatures (Handel et al. 2014). In our study, the presence of unsuitable habitats for one species (*S. texanum*) may necessitate spatial niche overlap in habitats that are suitable. Such overlap is typical to spatially structured ecosystems with scarce resources (savanna grasses: Kulmatiski et al 2020; carabid species Tsafack et al 2021). Together, our study showed that while large scale spatio-environmental niche partitioning did contribute to species favoring certain sites and excluding others, ultimately, it alone proved insufficient to comprehensively explain species coexistence.

### Finer spatiotemporal scales: Within-site coexistence via aggregation

Despite the differences in environmental preferences, nematodes co-occurred within sites and even within soil samples (Fig. 4A). Resource partitioning can be weak between species belonging to the same genus because of their preference for the same habitats or host species (Takahashi *et al* 2005). However, if suitable habitats are distributed discretely in space, then aggregation across habitat patches can enable coexistence on a single resource type, if intraspecific aggregation is greater than interspecific aggregation (Ives 1991, Roberts & Dobson 1995). Similar to demonstrations in insect communities (Takahashi et al 2005; Jaenike & James 1991), we found that aggregation can promote EPN species coexistence at the within-site scale. Nematode distributions within each site were aggregated (Fig. 4B), most likely due to the limited availability of insect hosts (Fig. 2A; Table S2). Both patterns that are consistent with previous work on EPNs (Stuart & Gaugler 1994; Spiridonov *et al* 2007; Půža & Mráček 2010). Additionally, our field patterns reveal that intraspecific aggregation each for species was greater than their interspecific aggregation, meeting the conditions for species persistence (Fig. 4B-D). If clumped dispersion offers a survival advantage to parasites, it is likely that their motile infective stages would naturally exhibit aggregation without any external stimuli. Shapiro-Ilan et al. (2014) discovered that several EPN species indeed exhibit high intraspecific aggregation patterns in homogenous arenas, even in the absence of host cues. Therefore, we show that aggregation across patches within a site can facilitate nematode species coexistence at this local scale.

Importantly, the relatively low interspecific aggregation we observed stems from random association between species (*C*_*x*,*y*_ ∼ 0) and not negative association (*C*_*x*,*y*_ < 0, see Figure 1 C i vs. iii). A negative association between species would suggest fine scale niche partitioning or a competition-colonization trade-off (Sevenster & van Alphen 1996). Instead within our field sites, nematodes exhibited random aggregation between species, a pattern often seen in communities with low host specialization (Ives 1991; Wertheim et al., 2001). For example, in both marine fish ectoparasites and rodent flea communities, it is intraspecific aggregation, not negative associations across species, that is predicted to enable parasite species coexistence (Morand et al 1999; Krasnov et al 2006). Thus, our work adds to a growing consensus that intraspecific aggregation, a well-known pattern of parasite distributions, may be a key feature predicting increased parasite diversity.

### Within-host fitness consequences of aggregation on parasite fitness

Aggregation had non-linear consequences on parasite fitness imposing negative brake on its own population growth. Besides demonstrating that the observed field patterns of aggregation support species coexistence, a lab mesocosm experiment evaluated the assumption that greater intraspecific aggregation intensifies intraspecific competition. This study revealed how heightened intraspecific aggregation can lead to competition among two EPN species (Fig. 5B). In both species, increasing number of infecting juveniles in a patch had a humped-shaped effect on total number offspring produced (Fig 5A). Specifically, number of offspring decreased when too few or too many parasites co-occurred in a patch and peaked at intermediate densities. An analogous pattern occurs when a single insect host is inoculated with a varying dose of nematodes (Selvan et al 1993). In parasitic nematodes, when too few parasites colonize a host, they have trouble clearing the immune system or exploiting host resources effectively (Keymer 1982). On the other hand, when too many parasites colonize an individual host, intraspecific competition for host resources reduces the yield per host. Aggregation implies that soil patches (or hosts) have either too few or too many parasites. The fitness effect of aggregation estimated for the two species at finer spatiotemporal scales (i.e., within-site scale) showed that increasing aggregation strongly reduced *per capita* fitness, dampening the growth of each species. Therefore, aggregation leads to a strong negative feedback which can stabilize nematode species coexistence in the field.

### Future direction

Mechanisms governing parasite species coexistence across spatiotemporal scales could be expanded in the future. First, explicit comparison of niche partitioning and spatial aggregation can partition the relative strength of each mechanism in maintaining species coexistence. Spatial aggregation among mycophagous *Drosophilid* species has been identified as more critical for species coexistence than resource partitioning (Wertheim et al 2000; Toda et al 1999), though their relative importance can vary based on the community’s phylogenetic structure (Takashi et al 2005). While resource partitioning plays a significant role in reducing competition among distantly related species, it appears less crucial for closely related species. In our study of four host-generalist species of nematodes from the same genus, we found a high degree of niche overlap at a local scale. Furthermore, our study identified few potential hosts among the sampled macroinvertebrates, accounting for less than 1% of the variation in nematode distributions. Yet the high degree of intraspecific aggregation we observed most likely reflects the tendency to of EPNs to emerge *en masse* from an insect carcass and move in a coordinated fashion (Shapiro-Ilan et al 2014). Future work could expand on the spatial or phylogenetic scope to reveal stronger environmental patterns. Implementing a more integrated sampling approach, such as month-long bait traps, could help quantify the role of resource partitioning in EPN communities more effectively.

Second, extending the study to multiple years could enhance our understanding of spatiotemporal parasite niche partitioning. Our one-year study indicated that the species exhibited more similar temporal usage patterns rather than distinct temporal profiles. However, what happens when conditions get more extreme and less predictable? How do species that are unfavoured by the environment sustain one bad year after another and not go extinct? A multiyear approach could better link these temporal patterns to variation mechanisms based on species-specific responses to environmental fluctuations (Chesson 2000). For example, if environmental infective stages respond differently to environmental fluctuations and have some mechanism to buffer them through unfavourable conditions, then species can coexist via the storage effect (Wisnoski & Lennon 2021). Such data may show that it is variation *per se* that is important for coexistence in this system.

Finally, within-host partitioning of resources like immune and energy niches may predict the effect of environmental aggregation on species coexistence (Ramesh & Hall, 2023). Theory suggests that aggregation tends to benefit species when increased coinfection leads to negative fitness outcomes (Morrill et al 2017), especially if parasites are large, site-specific, or trigger cross-immunity (Lello et al 2004; Bush & Malenke 2008). Aggregation may increase with higher site specificity and overlap between species, as it reduces interspecies encounters, or decrease when parasites have lower site specificity and are less prone to direct resource competition. Future experiments can mechanistically link how such join exploitative and immune-mediated apparent competition via manipulation of say immune-suppression can shape aggregation (Johnson & Hoverman 2014). Together, these expansions of between-host parasite dynamics can produce insights into coexistence, stability of food webs in ecosystems or even human infectious diseases.

## Conclusion

The maintenance of parasite species diversity requires due consideration of the environmental distributions of parasite infective stages even prior to host encounter. To test competing mechanisms of species coexistence, we borrowed ideas from old and new niche theories of species coexistence stemming from community ecology and applied it to the parasite realm. Using year-long field sampling of entomopathogenic nematodes, we demonstrate that species can coexist across multiple scales. At a larger spatiotemporal scale environment niche partitioning can favour species in some sites and exclude them in others. At finer spatiotemporal scales, intraspecific spatial aggregation can lead to coexistence through negative feedbacks of each species on its own population growth. While our work capitalized a parasite taxon with a charismatic transmission stage, persistence and environmental aggregation in this stage is a key component affecting disease dynamics in most parasite systems (Dobson & Roberts 1994). Together, we assert that adopting a niche-based perspective to link environmental tolerances and distributions of parasite transmission stages can only enhance predictive insight into maintenance of parasite diversity and its link to ecosystem functioning alike

## ACKNOWLEDGMENTS

AR was supported by the Graduate Women in Science National Fellowship (GWIS) and Indiana University (IU) Center for the Study of Integrative Animal Behavior (CISAB) Fellowship. This work was supported by an Indiana Academy of Science Senior Grant Award and an IU Research and Teaching Preserve (IURTP) Research Awards. We gratefully acknowledge NSF DEB 0919015 and 0515832 and C Lively for supporting foundational work on this system. OMS was supported by the Emerging Scholars REU, IU Center of Excellence for Women & Technology. We thank E Huenupi and K Beidler for sharing valuable knowledge about soil sampling; M Chitwood for maintaining the IURTP properties. We are grateful to L Whitney, L Totaro, and especially RK Phillips, SN Ramesh, and S Ramesh for their assistance in the field and in the laboratory even as we sampled through a global pandemic. AR credits B Joel and MD Pallavi for artistic inspiration.

## Data accessibility statement

All data files and associated R code will be published in a public repository upon acceptance

## SUPPLEMENTAL APPENDIX

In this supplemental appendix, we provide additional support to underlie our methods and results in the main text.

Section 1 (S1): describes the calculations for nematode abundance from DNA extraction and quantification using real-time qPCR.

Section 2 (S2): provides supporting information to explain the limited niche partitioning observed at the large temporal and spatial scales *i*.*e*. between-seasons (temporal partitioning; Fig. 2) and between-sites (spatial partitioning; Fig. 3).

Section 3 (S3): provides additional information to explain niche partitioning at the finer spatial scale *i*.*e* within-site scale via aggregation (Fig. 4) and how fitness consequences of aggregation facilitates coexistence in the nematode community (Figs. 5).

All statistics and visualizations were using R (version 4.0.5) (R Core Team, 2020).

### Section 1 (S1) Quantifying nematode abundance

#### S1.1. Estimating nematode abundance: DNA extraction and nematode quantification using real-time qPCR

Nematodes were extracted from the soil by sucrose centrifugation after soil samples were sieved using 2000 and 20uM meshes separates the nematode as described in the main text (following Jenkins 1964, modified by Jaffee, B at UCSD). After centrifugation, the supernatent was poured through the sieve combination, rinsed and then gathered in to a 1.5 ml microcentrifuge tube with no more than 0.5 ml of remaining sediment. The samples were stored at 4 ^0^C. DNA was extracted from the frozen samples using the manufacturers protocols for the maximum yield using the Qiagen DNeasy PowerSoil Pro Kit. The DNA was resuspended in 100 mL of elution buffer and stored in 4 ^0^C until use. All DNA samples were analysed using a quantitative PCR (qPCR) approach.

Species-specific primers and probes were designed from the ITS and 28S region (Campos-Herrera et al. 2011b) for all the EPNs using sequences from the target native species (Table S1). Real-time PCR quantification was performed for each sample using primer from each of the four EPN species, following protocols described by Sinkiewicz *et al* 2016 (Centre for Integrative Animal Behaviour lab, Indiana University). Briefly, the reactions were performed in a final volume of 25 uL, using 12.5 uL of the SYBR Green Master Mix (Applied Biosystem, USA), 1.5 uL of forward and reverse primer combination for each species, and 3mL of the DNA sample. Reactions were performed in optical 96-well reaction plates on LightCycler 480 System (Roche Diagnostics) with thermal cycling for the nematode. Thermal cycling was performed under the following conditions: 10 minutes of pre-incubation at 95°C, followed by 45 cycles of amplification 95°C for 30 s, 55°C for 1 min and 72°C for 1 minute, followed by 1 cycle of 60°C for 1 minute during melting phase, and finally 1 cycle of cooling at 40°C for 1 minute. All samples were run in duplicates. The mean of the threshold cycle values (Ct) corresponding to each of the four species was recorded for each sample. Then, the Ct values were analysed against a standard curve for each sample to quantify corresponding number of juveniles of each species in a given sample.

The standard curves for all species were performed separately with the same Qiagen DNeasy PowerSoil Pro kit. DNA was extracted from multiple replicate samples of known quantities of nematodes for each species using qPCR approach as described above. Linear regressions of natural log (ln) of number of nematodes and ln threshold cycle value (Ct) were performed to derive standard curves for each nematode species (Fig. S1). Then, the estimated slope and intercept were used to calculate number of infective juveniles of each species in field samples. Based on the machine recommendation to disregard *Ct* values at or above 40, and the standard curves, our detection level was 1 nematode for *S. affine, S. costaricense*, and *S. texanum* and 4 nematodes for *S. kraussei*.

**Table S1:**
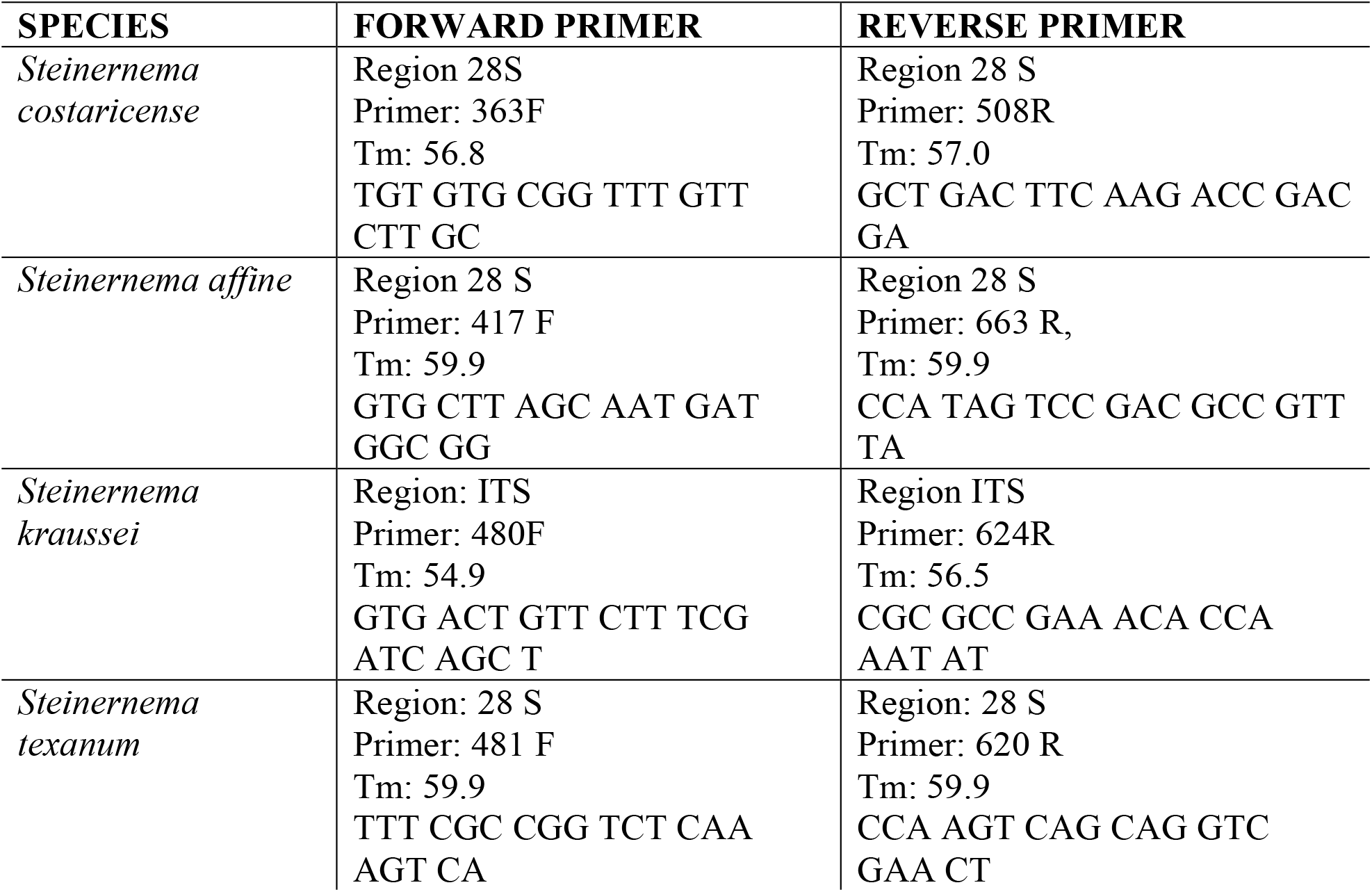
Species-specific quantitative PCR primer designs for four sympatric EPN species.

**Fig. S1.**
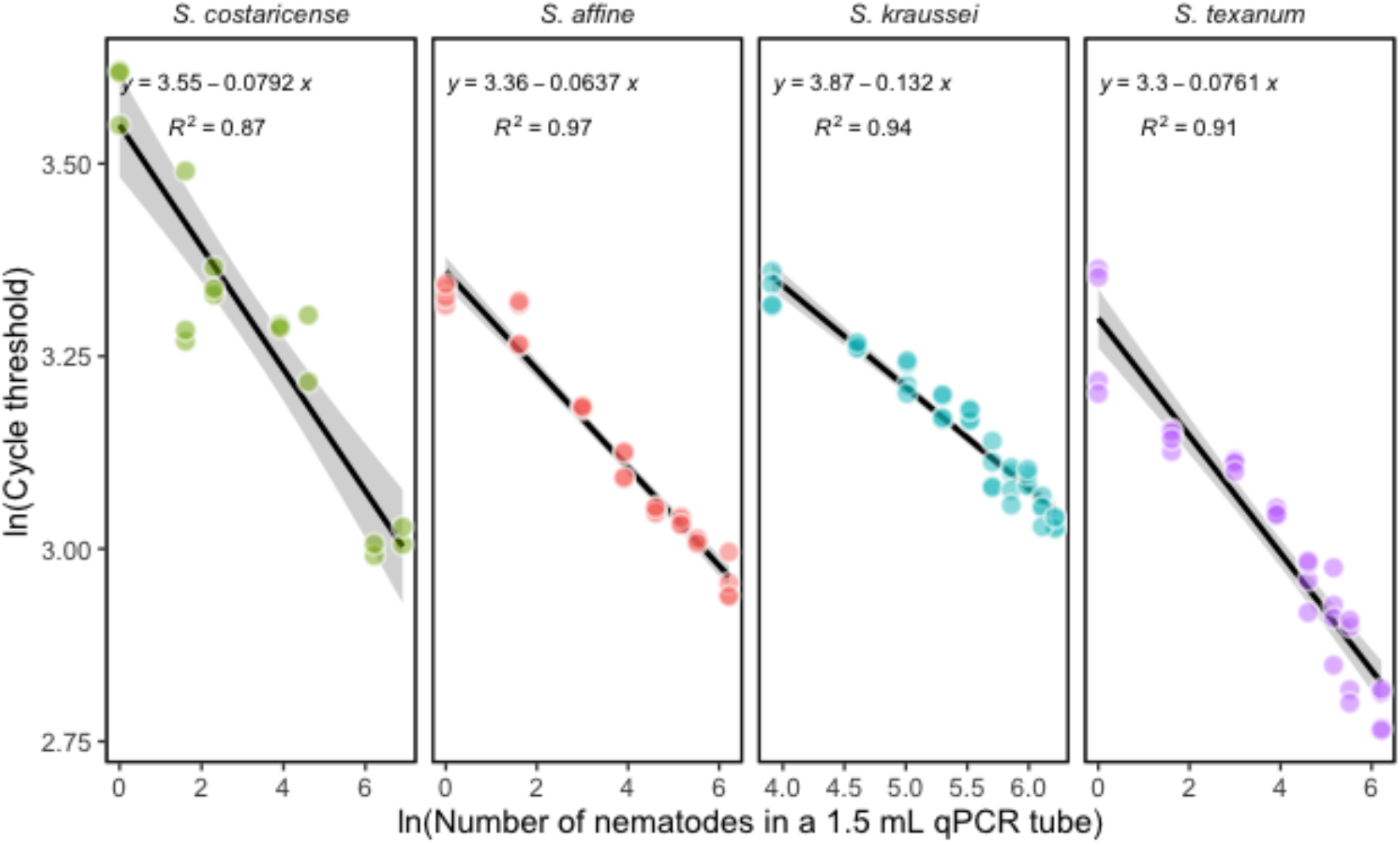
Standard curves calculated from linear regression of log threshold cycle value (Ct) and number of infective juveniles for four EPN species. Inset equations: slope, intercept, *R*^*2*^, and error ribbons (gray) indicate 95% confidence intervals.

**Table S2:**
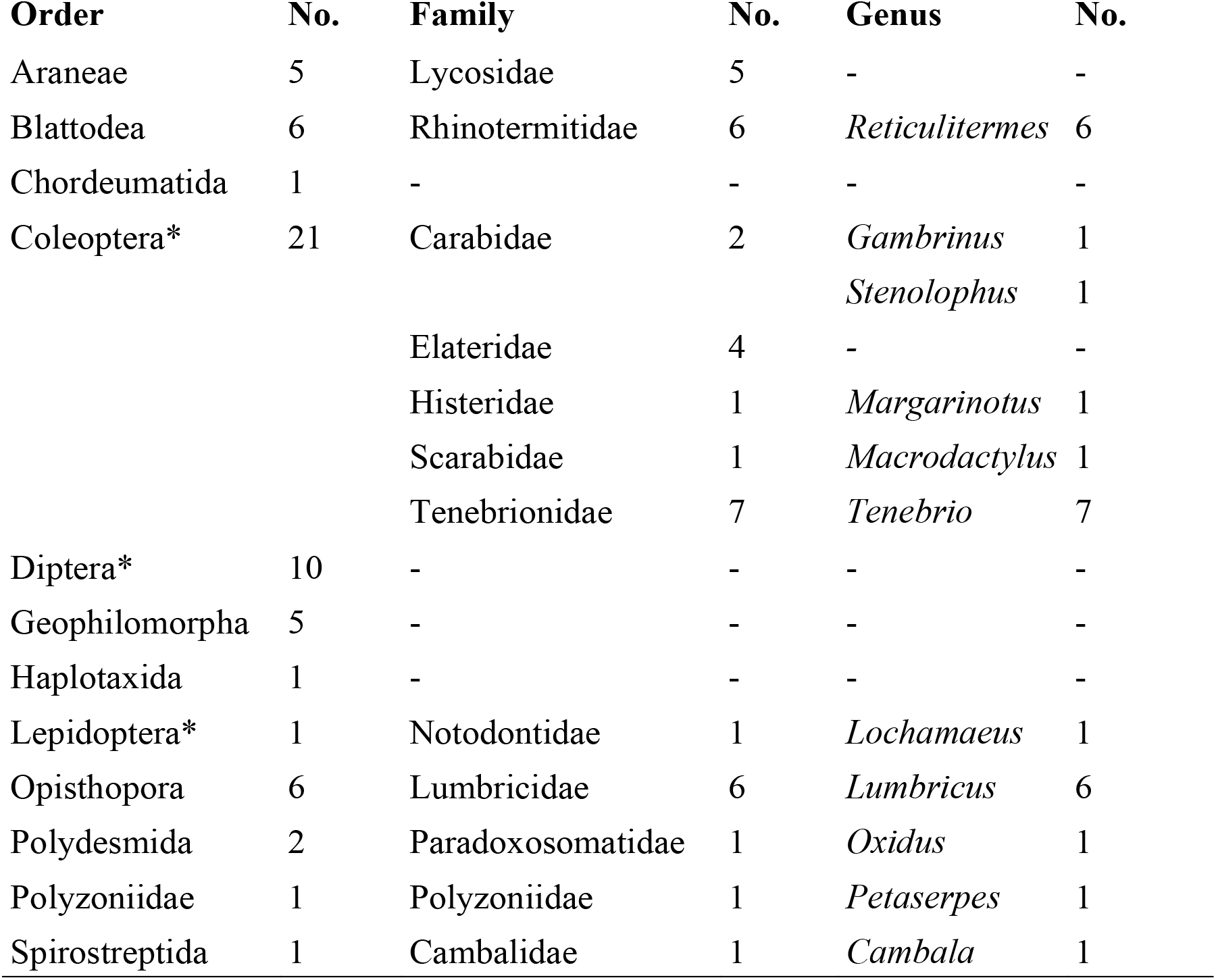
Macroinvertebrates identified in soil samples. Each individual larger than (3 mm) was identified to order, family or genus as possible. Potential EPN hosts marked with *, typically soft-bodied insect larvae.

### Section 2 (S2) Large scale temporal and spatial niche partitioning

#### S2.1 Niche overlap index

We calculated the niche overlap index at two major spatio-temporal scales. First, at the temporal scale we calculated the overall extent of temporal niche overlap pooled for each site and across each month. Then, we calculated the extent of temporal overlap at each of the six sites. Second, at the larger spatial scale (across sites), we calculated the overall extent of spatial niche overlap pooled across sites and seasons. Then, we calculated the extent of spatial overlap for each month. The niche overlap index at both scales indicate that species do not exhibit strong niche partitioning along the temporal or spatial axis.

**Table S3:**
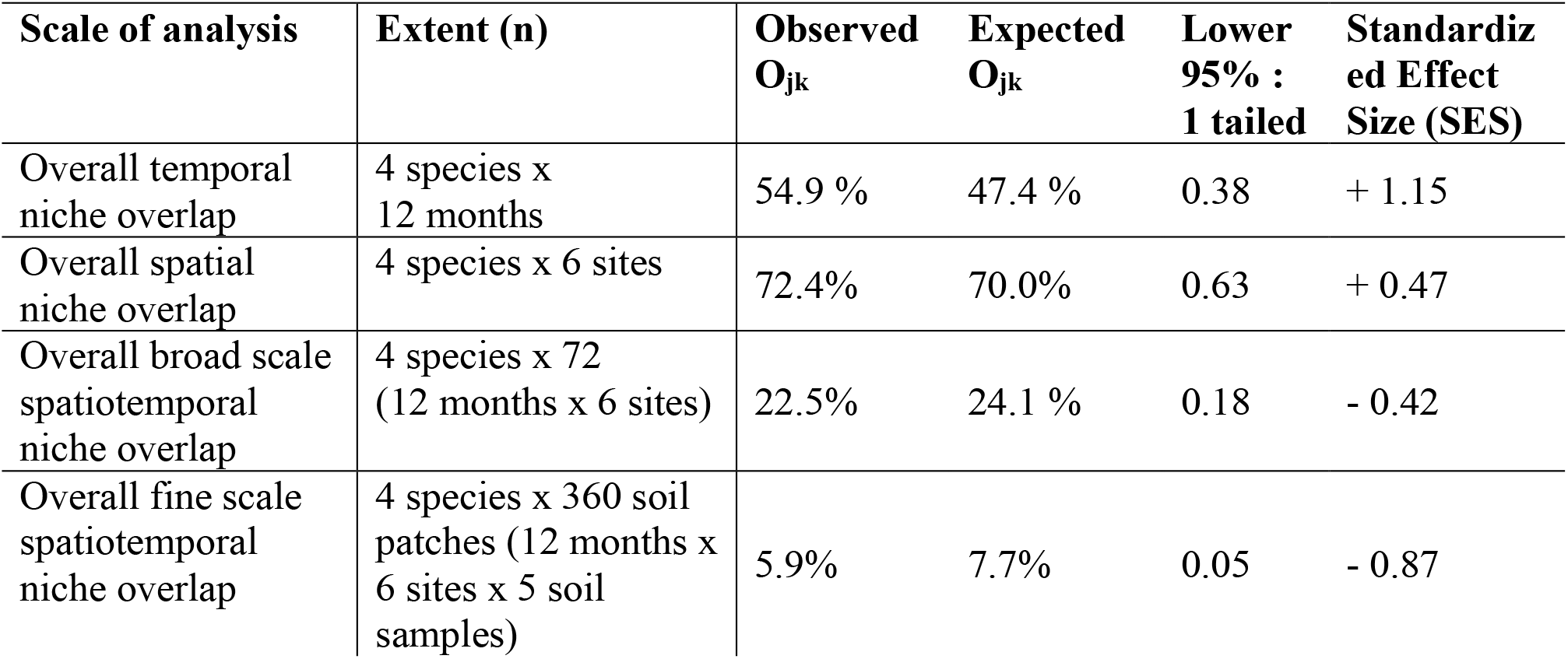
Pianka’s niche overlap index for large temporal (between months) and spatial (between sites) scales using the RA4 index. A one-tailed lower 95% confidence interval tests for significant niche partitioning between observed and expected niche overlap (O_jk_). The Standardized Effect Size (SES) converts the *p*-value into a standardized deviate. Large positive values of the SES indicate increasingly small upper-tail probabilities, and large negative values of SES indicate increasingly small lower-tail probabilities. Non-significant tail probabilities usually fall between −2.0 and +2.0

| Scale of analysis | Extent (n) | Observed<br>$O_{jk}$ | Expected<br>$O_{jk}$ | Lower<br>95% :<br>1 tailed | Standardiz<br>ed Effect<br>Size (SES) |
| --- | --- | --- | --- | --- | --- |
| Overall temporal<br>niche overlap | 4 species x<br>12 months | 54.9 % | 47.4 % | 0.38 | + 1.15 |
| Overall spatial<br>niche overlap | 4 species x 6 sites | 72.4% | 70.0% | 0.63 | + 0.47 |
| Overall broad scale<br>spatiotemporal<br>niche overlap | 4 species x 72<br>(12 months x 6 sites) | 22.5% | 24.1 % | 0.18 | - 0.42 |
| Overall fine scale<br>spatiotemporal<br>niche overlap | 4 species x 360 soil<br>patches (12 months x<br>6 sites x 5 soil<br>samples) | 5.9% | 7.7% | 0.05 | - 0.87 |

**Fig. S2.**
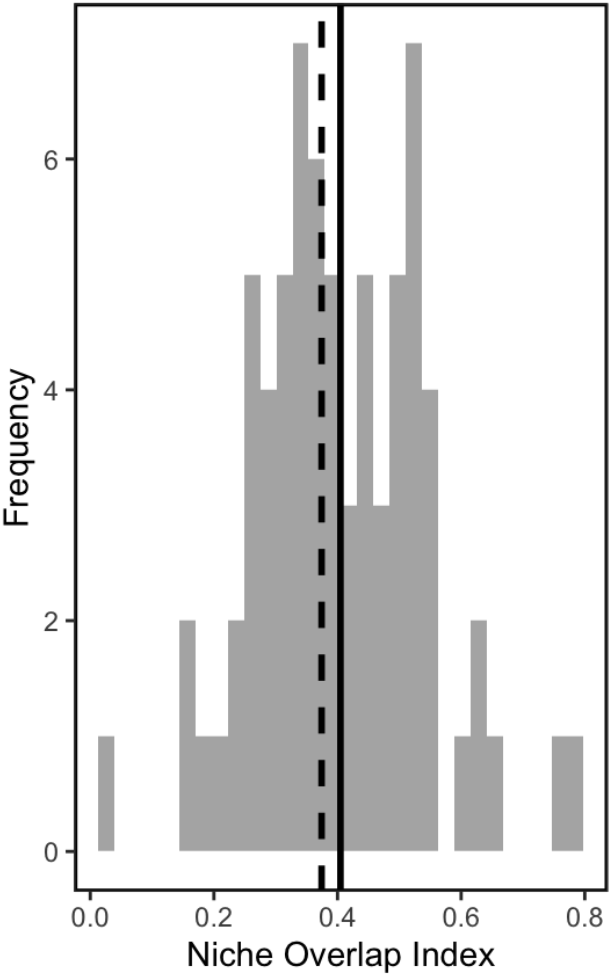
Fine scale spatiotemporal niche overlap by site month-combination: A frequency distribution of species niche overlap index calculated within each site each month suggests that species overlap frequently (with only 2 of 72 site/ month combinations showing niche partitioning). Overall, the mean of the all the observed niche overlap index (black, solid) was 0.40 and simulated was 0.37 (black, dashed)

#### S2.2 Spatiotemporal dynamics of nematode community

To examine how nematode abundances varied across site-month combinations we used a multivariate analysis. Here we present the statistics for a RDA with a full model including all the environmental variables for this dataset and generated both the overall contribution and percent contribution of each environmental variables (*R*^*2*^_*adj*_; Table S4). Since niche elevation accounted for the most variation in species abundance, we conducted a mixed-model ANOVA. This analysis tested how species abundance, soil temperature, moisture, and pH varied across six sites: two in each of three elevation categories (Highlands at 218 m, Midlands at 216-212 m, and Lowlands at 205-208 m), with each site 60 meters apart. The total number of observations was 72 (6 sites × 12 months).

**Table S4:**
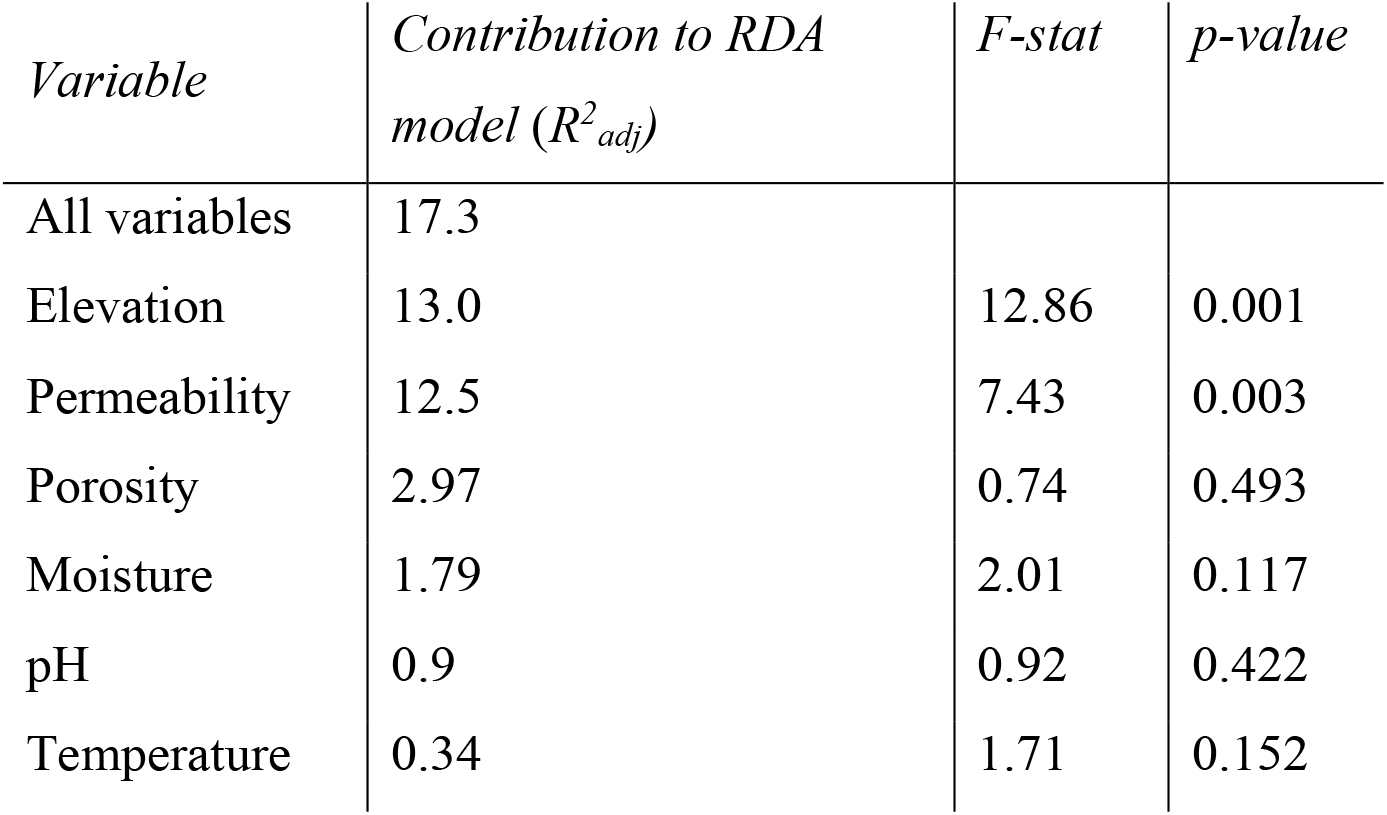
The environmental variables and their percent contribution (*R*^*2*^_*adj*_) to nematode variation.

**Table S5A:** Soil properties (mean ± SE) of the six sites at the Moores Creek Preserve along an elevational gradient across six study sites, with two sites each in (Highland ●, Midlands ▲ and Lowlands ◼) as shown in Fig. 3C-E.

| *Variables* | *Highland* |  | *Midland* |  | *Lowland* |  | *F-stat* | *df* | *p-value* |
| --- | --- | --- | --- | --- | --- | --- | --- | --- | --- |
|  | *Site 1* | *Site 2* | *Site 3* | *Site 4* | *Site 5* | *Site 6* |  |  |  |
| Elevation (m) | 218 | 218 | 216 | 212 | 208 | 205 |  |  |  |
| Soil moisture (%) | 14.7<br>$\pm 2.5$ | 15.4<br>$\pm 2.2$ | 15.1<br>$\pm 2.2$ | 16.2<br>$\pm 2.5$ | 22.2<br>$\pm 2.3$ | 26.6<br>$\pm 2.0$ | 32.3 | 5, 55 | <0.01 |
| Soil temperature ($^{\circ}$C) | 14.8<br>$\pm 2.1$ | 14.7<br>$\pm 2.1$ | 15.0<br>$\pm 2.1$ | 15.2<br>$\pm 2.1$ | 15.2<br>$\pm 2.1$ | 14.6<br>$\pm 2.1$ | 10.7 | 5, 55 | <0.01 |
| pH | 4.6<br>$\pm 0.1$ | 4.5<br>$\pm 0.1$ | 4.6<br>$\pm 0.1$ | 4.5<br>$\pm 0.1$ | 4.8<br>$\pm 0.1$ | 4.7<br>$\pm 0.1$ | 3.3 | 5, 55 | <0.01 |
| Soil porosity | 69.2<br>$\pm 2$ | 61.4<br>$\pm 0.9$ | 48.4<br>$\pm 3.3$ | 58.4<br>$\pm 2.6$ | 49.6<br>$\pm 1.3$ | 53.0<br>$\pm 1.6$ | 14.0 | 5, 66 | <0.05 |
| Soil permeability (minutes) | 2.9<br>$\pm 0.1$ | 3.0<br>$\pm 0.0$ | 3.1<br>$\pm 0.1$ | 2.7<br>$\pm 0.1$ | 2.0<br>$\pm 0.0$ | 2.0<br>$\pm 0.0$ | 62.6 | 5, 55 | <0.01 |

**Table S5B:**
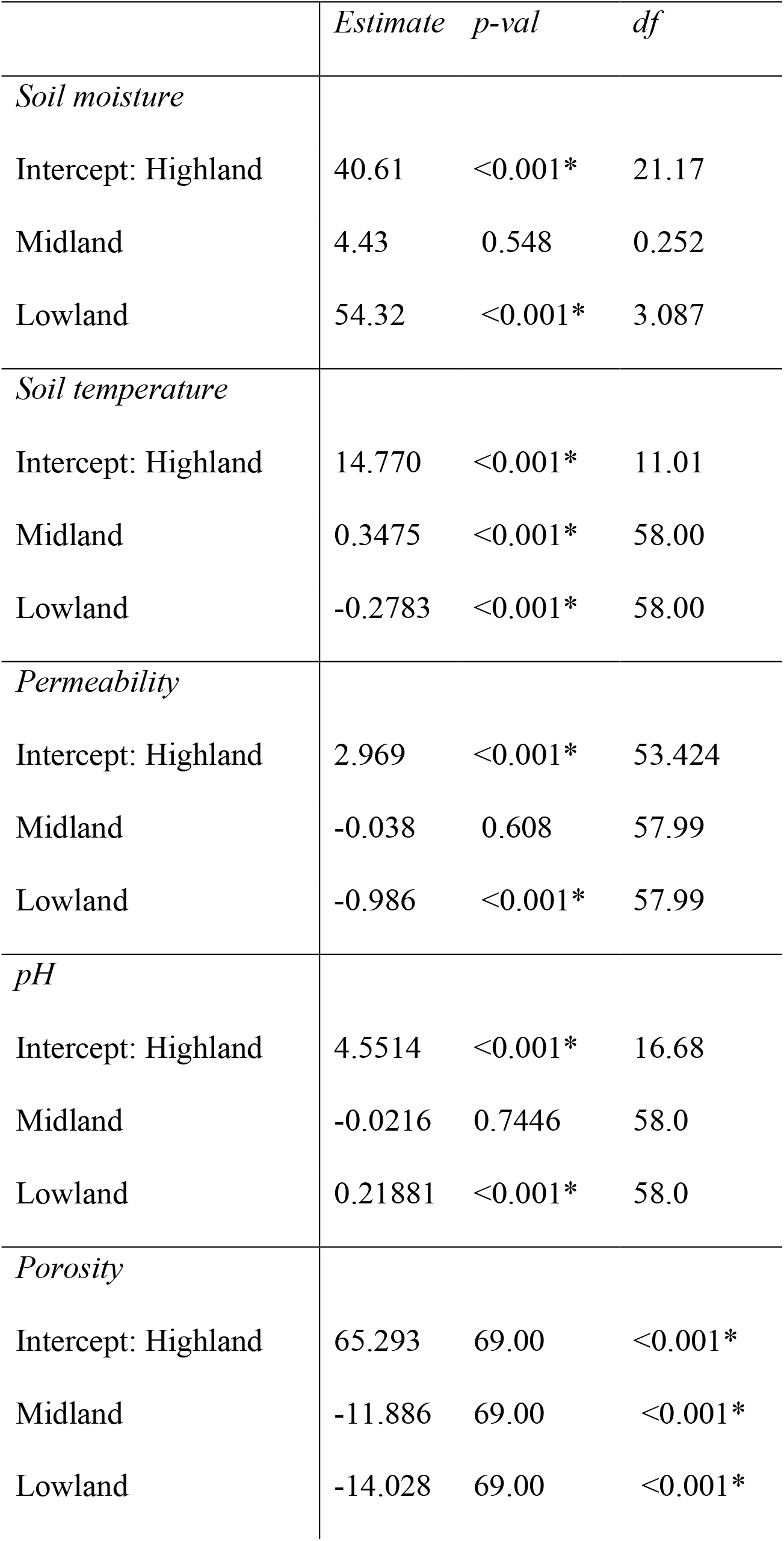
Table S5B. Shifts in climate variables: soil moisture (%) and temperature (^0^C), and soil variables: permeability (min), pH, and porosity along the altitudinal gradient. These relationships were tested using generalized linear models with month as random effects with gaussian for all soil variables.

**Table S5C:** Shifts in species abundance along the altitudinal gradient with two sites each in (Highland ●, Midlands ▲ and Lowlands ◼) as shown in Fig. 3C-E. These relationships were tested using generalized linear models with month as random effects with negative binomial family for species abundances.

|  | *Estimate* | *p-val* | *F* | *df* |
| --- | --- | --- | --- | --- |
| *Species abundance* |  |  |  |  |
| *S. affine* |  |  |  |  |
| Intercept: Highland | 4.94 | <0.001* | 2.58 | 2, 67 |
| Midland | 0.23 | 0.653 |  |  |
| Lowland | -1.00 | 0.052 * |  |  |
| *S. costaricense* |  |  |  |  |
| Intercept: Highland | 3.35 | <0.001* | 8.0 | 2, 67 |
| Midland | 0.08 | 0.716 |  |  |
| Lowland | 0.92 | <0.001* |  |  |
| *S. kraussei* |  |  |  |  |
| Intercept: Highland | 4.06 | <0.001* | 3.08 | 2, 67 |
| Midland | -0.33 | 0.338 |  |  |
| Lowland | -0.88 | <0.005* |  |  |
| *S. texanum* |  |  |  |  |
| Intercept: Highland | 3.00 | <0.001* | 904 | 2, 67 |
| Midland | -1.86 | <0.01* |  |  |
| Lowland | -5.94 | <0.001* |  |  |

### Section 3 (S3) Fine scale niche partitioning via spatial aggregation

#### S3.1. Species persistence via aggregation

Species persistence via aggregation requires three key indices measuring the intra- and inter-specific aggregation.

##### Intraspecific aggregation

*J*_*x*_ is the index of intra-specific aggregation where if *J*_*x*_ = 0, then variance is equal to then mean and species are randomly distributed, *J*_*x*_ > 0 implies variance much greater than the mean *i*.*e*. over-dispersion or aggregation, and *J*_*x*_ < 0 is uniform distribution. For a community with two species *x* and *y*, intraspecific aggregation *J* is measured for both species.

##### Interspecific aggregation

*C*_*x*,*y*_ is the index of inter-specific aggregation between the two species where *C*_*x*,_ = 0, then there is random association between the species. If *C*_*x*,_ > 0, there is positive association implying high potential for interspecific competition. If *C*_*x*,_ < 0 there is negative association (or no overlap) indicating resource segregation and no potential for interspecific competition within a patch.

##### Index of Persistence

Together these indices indicate species persistence (Sevenster 1996). Species persistence is measured by the Index of Persistence (*A*_*x*,*y*_) where,

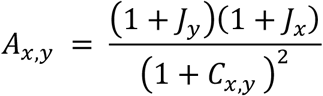

Here, if the joint product of intraspecific aggregation of the two species is greater than their interspecific aggregation (*i*.*e. J*_*y*_ > *C*_*x*,*y*_) then species persist i.e., *A*_*x*,*y*_ > 1.

In summary, all the focal species were sufficiently aggregated (*J* > 0) where *S. costaricense* showed the lowest aggregation. The interspecific competition *C*_*x*,*y*_ was centered around zero indicated random association. Taken together, the index of persistence *A*_*x*,*y*_, predicts that overall each species when present together with its competitors predicts persistence. Here, we log-transformed *A*_*x*,*y*_ and applied the glm.nb function from the MASS package (Venables & Ripley 2002) to implement a generalized linear model with a negative binomial distribution. This allowed us to test whether *A*_*x*,*y*_ > 1 or *ln*(*A*_*x*,*y*_ + 1) > *ln*(2) = 0.69. We then used the 95% confidence intervals to determine if the estimates fell within the range of their respective thresholds

**Table S6.**
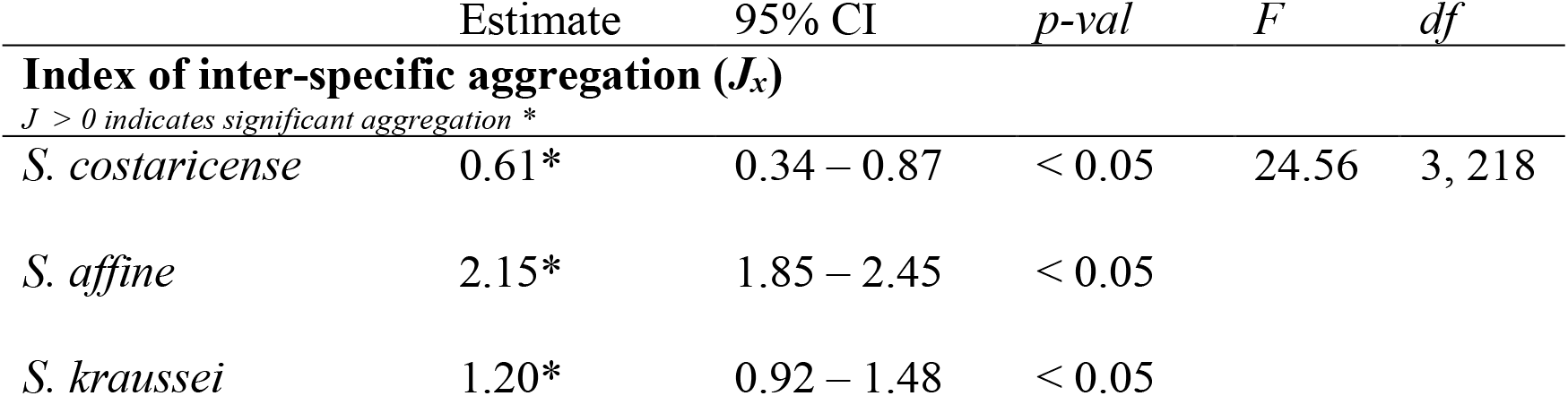
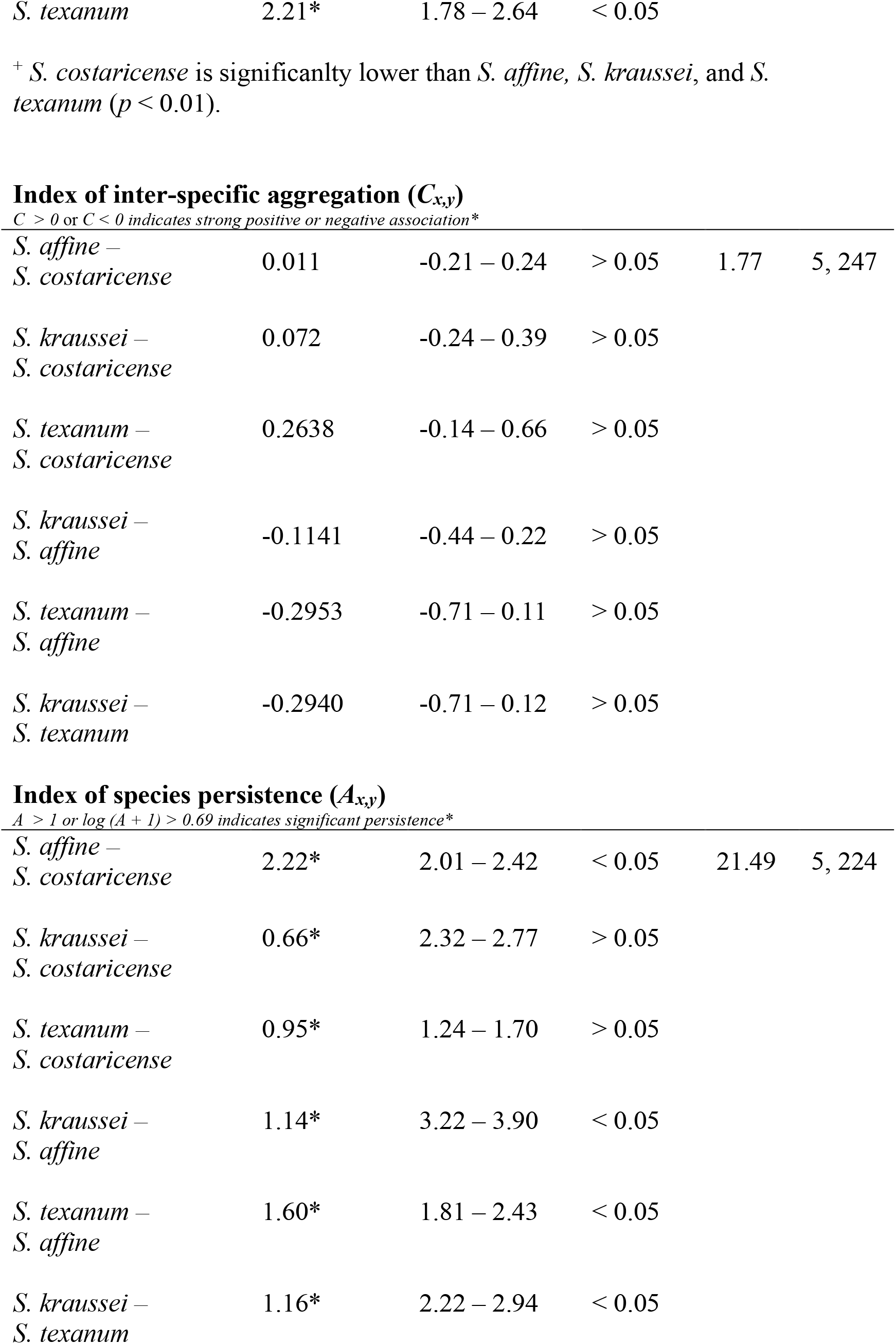
Linear mixed model fits with site and time as random factors i.e. estimates, 95% confidence intervals for each estimate for indices of aggregation: intraspecific self-aggregation (*J*), interspecific aggregation (*C*_*x*,*y*_) and their joint effect on species persistence (*A*_*x*,*y*_) for all four EPN species pairs.

|  | Estimate | 95% CI | *p-val* | *F* | *df* |
| --- | --- | --- | --- | --- | --- |
| **Index of inter-specific aggregation ($J_x$)** |  |  |  |  |  |
| *J > 0 indicates significant aggregation \** |  |  |  |  |  |
| *S. costaricense* | 0.61* | 0.34 – 0.87 | < 0.05 | 24.56 | 3, 218 |
| *S. affine* | 2.15* | 1.85 – 2.45 | < 0.05 |  |  |
| *S. kraussei* | 1.20* | 0.92 – 1.48 | < 0.05 |  |  |
| *S. affine* –<br>*S. costaricense* | 0.011 | -0.21 – 0.24 | > 0.05 | 1.77 | 5, 247 |
| *S. kraussei* –<br>*S. costaricense* | 0.072 | -0.24 – 0.39 | > 0.05 |  |  |
| *S. texanum* –<br>*S. costaricense* | 0.2638 | -0.14 – 0.66 | > 0.05 |  |  |
| *S. kraussei* –<br>*S. affine* | -0.1141 | -0.44 – 0.22 | > 0.05 |  |  |
| *S. texanum* –<br>*S. affine* | -0.2953 | -0.71 – 0.11 | > 0.05 |  |  |
| *S. kraussei* –<br>*S. texanum* | -0.2940 | -0.71 – 0.12 | > 0.05 |  |  |
| *S. affine* –<br>*S. costaricense* | 2.22* | 2.01 – 2.42 | < 0.05 | 21.49 | 5, 224 |
| *S. kraussei* –<br>*S. costaricense* | 0.66* | 2.32 – 2.77 | > 0.05 |  |  |
| *S. texanum* –<br>*S. costaricense* | 0.95* | 1.24 – 1.70 | > 0.05 |  |  |
| *S. kraussei* –<br>*S. affine* | 1.14* | 3.22 – 3.90 | < 0.05 |  |  |
| *S. texanum* –<br>*S. affine* | 1.60* | 1.81 – 2.43 | < 0.05 |  |  |
| *S. kraussei* –<br>*S. texanum* | 1.16* | 2.22 – 2.94 | < 0.05 |  |  |

### S3.2 Does intraspecific aggregation translate to intraspecific competition?

First, we estimate the nematodes densities within a soil sample in the field using the same sieving assay used to isolate the nematodes. Soil-filled garden pots were seeded with seven initial nematode density of *S. costaricense*: 0, 250, 500, 1000, 2500, 5000, 10000.

Approximately 300cm^3^ of soil, similar to the amount sampled in the field using a soil corer, was added to each pot. The nematodes were then sieved on 20uM sieves, centrifuged, and their abundance was estimated using qPCR standard curves (S1). We used a linear model to examine the relationship between initial and final nematode densities post-processing (*F*_*1*,*20*_ = 33.57, *p* < 0.001, *R*^*2*^ = 0.626). Using this relationship, we back-transformed the qPCR-quantified nematode quantities from the field samples (Fig. 4A) to estimate nematode abundance in the soil. This approach was applied specifically to estimate the abundance of *S. costaricense* and *S. kraussei*, as they have comparable body sizes and can be captured similarly on the sieve. The estimated nematode counts in each soil sample were utilized to determine the index of aggregation (*J*). Both species demonstrated aggregation patterns, with *S. costaricense* exhibiting lower aggregation compared to *S. kraussei*, consistent with the predictions made by the qPCR-only abundance data (Fig. 4A). To further validate the comparability of *J* between the two methods, we observed that the index of aggregation calculated using qPCR-only abundance data and transformed soil data displayed a 1:1 relationship. This provides additional confirmation of the consistency between the two approaches in assessing the aggregation behavior of the nematode species. Second, we estimated field fitness for each site based on lab density-fitness relationship. Our focus remained on the two species, *S. costaricense and S. kraussei*. Soil-filled garden pots were prepared with varying initial nematode densities (0, 100, 500, 1000, 5000, 10000) and one, three, or six *Galleria mellonella* larvae as hosts. We measured the total number of offspring emerging per host. The relationship between the number of infecting juveniles in the soil and the total number of offspring per patch was governed by a family of quadratic curves, observed across all combinations of host densities in both species (Fig. 5A-B). Specifically, too few or too many infecting nematodes resulted in a reduction in the number of offspring, while intermediate densities facilitated higher offspring numbers. Using the quadratic relationship for the case of a single host, we calculated the per capita nematode fitness *i*.*e*. 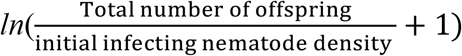 with increasing infecting nematode density *ln*(initial infecting nematode density) using a linear model (For a single host, *S. costaricense: F*_*1*,*15*_ = 109, *p* < 0.05, *R*^*2*^ = 0.871; *S. kraussei: F*_*1*,*11*_ = 16.22, *p* < 0.05, *R*^*2*^ = 0.559). Then, we calculated the estimated *per capita* fitness for each soil sample for each site-month combination (*n*_*max*_ = 5 soil patches x 6 sites x 12 months = 360). Finally, we examined how within site-level fitness related to intraspecific aggregation (*J*). The within-site level fitness was calculated by taking the mean of *per capita* across soil samples which was then regressed against the index of aggregation (*J*). Increasing aggregation reduced *per capita fitness* per site for both species (*S. costaricense*: *R*^2^ = 0.12, *p* < 0.05; *S. kraussei*: *R*^2^ = 0.48, *p* < 0.05; Fig. 5C). This suggests that the key assumption of the aggregation framework, that that greater intraspecific aggregation leads to increased intraspecific competition, is true for the two species of *Steinernema spp*.

**Fig. S3.**
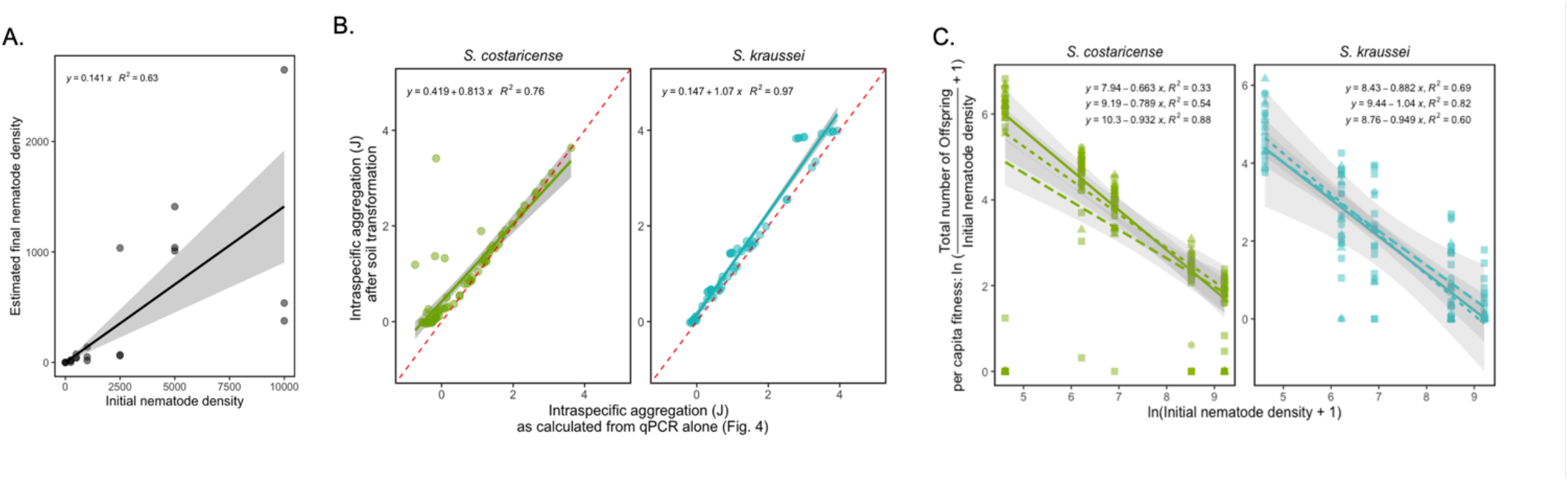
(A) Estimated the relationship between the intitial nematode density and final nematode density post seiving processing that were utilized to determine the index of aggregation (*J*). Both species demonstrated aggregation patterns, with *S. costaricense* exhibiting lower aggregation compared to *S. kraussei*, consistent with the predictions made by the qPCR-only abundance data (B) To further validate the comparability of *J* between the two methods, we observed that the index of aggregation calculated using qPCR-only abundance data and transformed soil data displayed a 1:1 relationship. (C) To estimated field fitness for each site based on lab density-fitness relationship, we calculated the *per capita* nematode fitness with increasing infecting nematode density presented for all three host densities here. The within-site level fitness was calculated by taking the mean of *per capita* across soil samples for one host case, which was then regressed against the index of aggregation (*J*). Increasing aggregation reduced *per capita fitness* per site for both species suggesting that aggregation facilitates coexistence in EPN communities.

**Table S7.** Linsear mixed models of total number of offspring regressed using quadratic fits of number of initial infecting nematodes for each host density treatment in *S. costaricense* and *S. kraussei* as shown in Fig. 5A: degree of freedom (*df*), F-ratio, *R*^*2*^_*adj*_ and p-value (* indicates statistical significance).

| Treatment | Linear<br>estimate | $p$-val | Quadratic<br>estimate | $p$-val | $df$ | $F$ | $R^2$ |
| --- | --- | --- | --- | --- | --- | --- | --- |
| *S. costaricense* |  |  |  |  |  |  |  |
| 1 host | 20.08 | <0.05 | -0.0014 | 0.055. | 2,18 | 12.63 | 0.53 |
| 3 host | 51.08 | <0.05 | -0.0057 | <0.05* | 2,21 | 13.68 | 0.52 |
| 6 host | 120.17 | <0.05 | -0.0094 | <0.05* | 2,21 | 15.73 | 0.56 |
| *S. kraussei* |  |  |  |  |  |  |  |
| 1 host | 9.86 | <0.05 | -0.0009 | <0.05* | 2,14 | 3.83 | 0.26 |
| 3 host | 6.45 | <0.05 | -0.0003 | 0.54 | 2,20 | 2.49 | 0.11 |
| 6 host | 31.14 | <0.05 | -0.0027 | <0.05* | 2,20 | 15.68 | 0.57 |

